# PAX2 is Transcriptionally Silenced by a Distinct Mechanism of Epigenetic Reprogramming to Initiate Endometrial Carcinogenesis

**DOI:** 10.1101/2024.12.23.630121

**Authors:** Subhransu S. Sahoo, Susmita G. Ramanand, Ileana C. Cuevas, Yunpeng Gao, Sora Lee, Ahmed Abbas, Xunzhi Zhang, Ashwani Kumar, Prasad Koduru, Sambit Roy, Russell R. Broaddus, Victoria L. Bae-Jump, Andrew B. Gladden, Jayanthi Lea, Elena Lucas, Chao Xing, Akio Kobayashi, Ram S. Mani, Diego H. Castrillon

## Abstract

In most cancers, including endometrial cancer, tumor suppressor genes harboring inactivating mutations have been systematically cataloged. However, locus-specific epigenetic alterations contributing to cancer initiation and progression remain only partly described, creating knowledge gaps about functionally significant tumor suppressors and underlying mechanisms associated with their inactivation. Here, we show that PAX2 is an endometrial tumor suppressor recurrently inactivated by a distinct epigenetic reprogramming event not associated with promoter hypermethylation. PAX2 is expressed throughout normal endometrial glands; however, microscopic clones with PAX2 protein loss arise spontaneously in adulthood and progress to endometrial precancers and cancers. Here we show that PAX2 protein loss occurs via transcriptional silencing in 80% of endometrial cancers. Mechanistically, transcriptomic, epigenomic, 3D genomic, and machine learning analyses showed that silencing is associated with replacement of open/active chromatin features (H3K27ac/H3K4me3) with inaccessible/repressive features (H3K27me3). The spread of the H3K27me3 signal is restrained by cohesin loops, preventing transcriptional dysregulation of neighboring genes. Functionally, PAX2 loss promoted endometrial carcinogenesis by rewiring the transcriptional landscape via global enhancer reprogramming. Genetically engineered mouse models together with organoid and human cell line studies established *Pax2* as an in vivo endometrial tumor suppressor that cooperates with *Pten* to produce lethal cancers. Our discovery of a specific and recurring epigenetic alteration that transcriptionally silences *PAX2* to initiate most endometrial cancers opens new lines of investigation into the origins of endometrial cancer with diverse implications for its diagnosis and treatment.

**One Sentence Summary:** We demonstrate that PAX2 is silenced by a highly recurrent epigenetic mechanism to initiate most endometrial cancers, establishing a novel mechanism of tumor suppressor inactivation.

## INTRODUCTION

Endometrial cancer (EC) accounts for 7% of all cancers and is the 4^th^ most common cancer in women, with ≥68,000 cases anticipated per year in the USA. In 2024, for the first time, EC deaths will exceed those of ovarian cancer (13,250 vs. 12,750) (*1*), underscoring EC’s significance as a women’s health issue. In contrast to declining incidence and improved survival for most cancers, EC incidence and mortality have been increasing over the past 40 years, by ∼1% each year (*1, 2*). Age is the most significant risk factor, with EC incidence peaking in the 7^th^ decade. Increasing life expectancy and other risk factors such as obesity contribute to EC’s rising incidence, but other poorly understood environmental and genetic factors are also at play (*3-5*).

EC arises from epithelial cells through a noninvasive histologic precursor termed endometrioid intraepithelial neoplasia (EIN), which progresses to invasive and lethal endometrioid adenocarcinoma (*6*). The EC landscape of somatically acquired driver mutations has been defined by next-generation sequencing (NGS) (*5-8*). *PTEN* is the most mutated gene (∼50% of EC) with *PIK3R1* and *PIK3CA* mutations also frequent, underscoring a central role of PI3K/PTEN signaling (*7*). Epigenetic transcriptional silencing through hypermethylation of a *MLH1* promoter CpG island is another mechanism of tumor suppressor inactivation in some EC (*9*). CpG methylation is detectable at the genomic level with single-base resolution by methylation NGS, which has established CpG island hypermethylation as a common mechanism underlying tumor suppressor inactivation (e.g., *APC*, *BRCA1*, *CDKN2A*, *MGMT, VHL*) (*10, 11*). However, NGS-based approaches may miss other types of non-mutational locus-specific epigenomic reprogramming events that may be critical or even initiating molecular driver events in cancer.

PAX2 is one of nine mammalian paired box DNA-binding transcription factors (TFs) (PAX1-9) with diverse roles in cell proliferation, lineage determination, organogenesis, and cancer. PAX2 is expressed in and is required for the development of the embryonic kidney and female reproductive tract; PAX2 knockout mice fail to develop kidneys or a uterus (*12*). Per reports in the clinical literature, PAX2 expression in endometrial glands persists into adulthood, but loss of PAX2 protein characterizes 80% of EIN and EC (*8, 13-17*). To date, there has been no adequate explanation of the mechanistic basis of PAX2 protein loss in the endometrium or its functional consequences.

Although it has been suggested that abnormal promoter methylation underlies PAX2 protein loss (*18*), this hypothesis lacks supporting evidence (*19*), and there have not been investigations establishing PAX2 as a functionally significant in vivo tumor suppressor. Here, we demonstrate through complementary approaches employing human specimens, cell lines, patient-derived xenografts (PDXs), and a conditional *Pax2* mouse EC model, that *PAX2* inactivation is an early (initiating) event caused by a specific non-mutational epigenetic reprogramming event unrelated to abnormal methylation but instead the replacement of open/active (H3K27ac and H3K4me3) with inaccessible/repressive (H3K27me3) chromatin features. These epigenetic processes occur within the confines of cohesin-mediated three-dimensional (3D) genomic architecture, thereby preventing transcriptional dysregulation of neighboring genes and limiting *PAX2* silencing as a focal epigenetic event. *PAX2* inactivation confers a competitive growth advantage driving endometrial cell outgrowth by reprogramming endometrial transcription via the commissioning and decommissioning of thousands of enhancers. These results establish *PAX2* epigenetic silencing as a specific, early, and highly recurring molecular driver event in EC, revealing a new paradigm for cancer-driving gene-level epigenetic alterations and opening new directions for EC research.

## RESULTS

### Emergence of PAX2-null clones in endometrium is age-dependent and PAX2 loss characterizes 80% of EC primary tumors and cell lines

PAX2 is expressed in Müllerian duct epithelium during embryogenesis (*12*), and strong expression persists in endometrial gland epithelium into adulthood, without expression in other uterine cell types (Fig. 1A). PAX2 protein loss has been reported in ≥80% of EC (Fig. 1B) (*8*). Minute PAX2-deficient clones (defined as loss of protein expression in all cells of ≥1 endometrial gland in cross-section) can be detected in some normal endometria, suggesting an early neoplastic event (*13*). To investigate the association of this phenomenon with age, we assessed PAX2 expression in endometria of women 18-25 and 44-45 y/o. These age groups were chosen because, in younger or older individuals, the endometrium is underdeveloped/atrophic due to low estrogen, making this the widest age range permitting meaningful assessments. PAX2 loss was not identified in 27 patients in the 18-25 y/o group. In contrast, clonal PAX2 loss was much more frequent (12/32 cases) in the 44-45 y/o group (Fig. 1C) (*8, 13, 14*). The difference between the age groups was statistically significant (Fig. 1D, P=0.00022), establishing that the emergence of PAX2-null clones is age dependent, as expected for a molecular event initiating endometrial neoplasia.

**Fig. 1.**
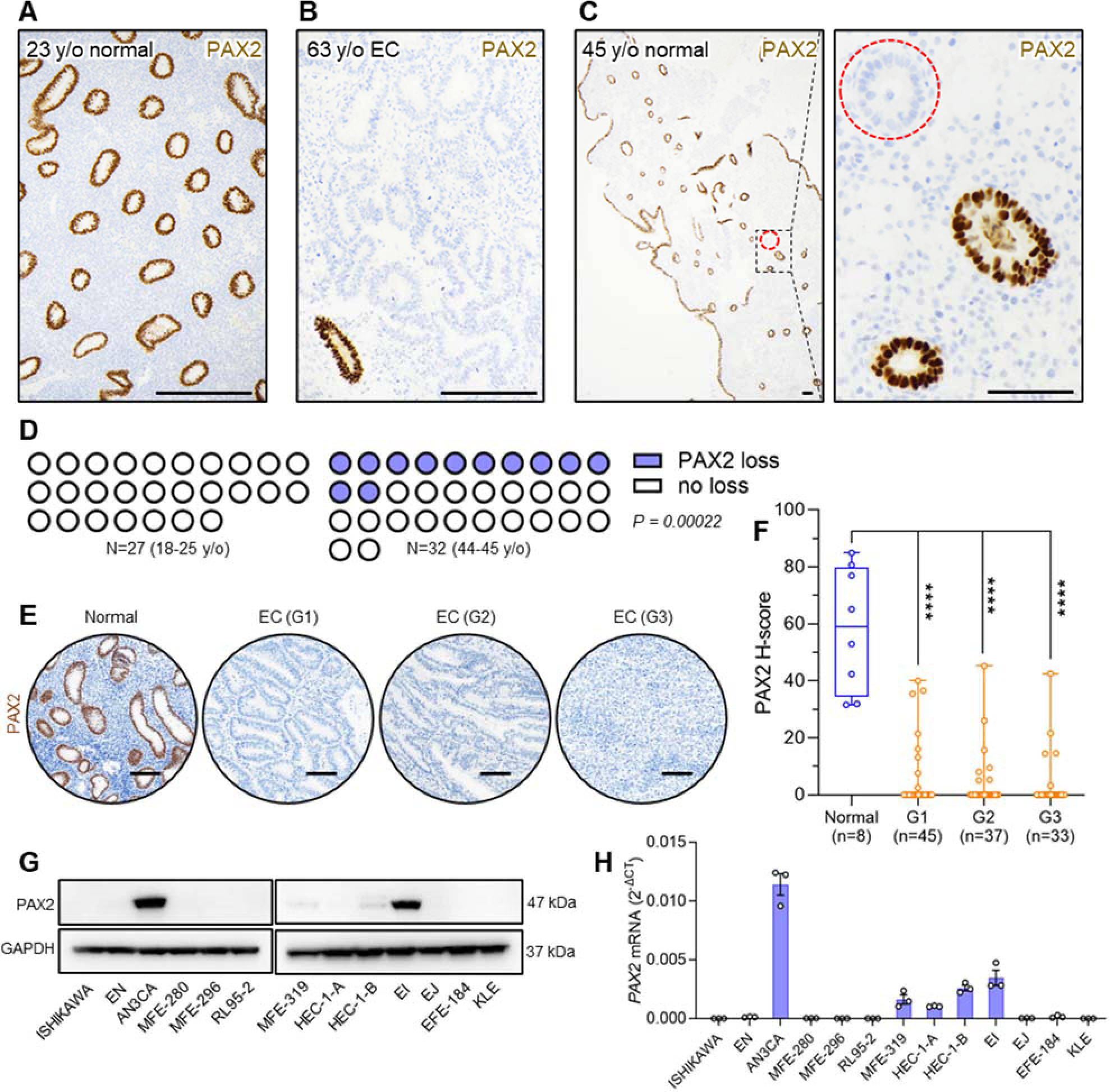
Emergence of PAX2-deficient clones in endometrial epithelium is age-dependent and associated with carcinogenesis. (A) Endometrial tissue section from younger (18-25 y/o) patient group, PAX2 immunolocalization. No PAX2-deficient clones were detected across entire specimen; representative region shown. Bar=200 μm. (B) EC from 63 y/o patient showing complete loss of PAX2, which occurs in 80% of EC. Residual normal (non-neoplastic) gland in lower left corner underscores striking and complete loss of PAX2 expression in EC. Bar=200 μm. (C) Endometrial tissue section from older (44-45 y/o) patient group. Dashed red circle highlights single gland in entire specimen with PAX2 loss; only portion of section shown. Right panel, magnification of boxed area showing complete (clonal) loss in all cells of the gland. Bars=100 μm. (D) Parts of whole plots show cases with PAX2 protein loss among younger (n=27) and older (n=32) patients. P value per 2-sided Fisher’s exact test. (E) PAX2 expression in normal proliferative endometrium and loss in most (>80%) ECs of grades 1-3. Bars=100 μm. (F) Box-and-whisker plots of PAX2 protein expression levels per H-scores in normal endometrium (n=8) and ECs (Grade 1, n=45; Grade 2, n=37; Grade 3, n=33). ****P<0.0001 per 2-tailed Mann-Whitney *U* test. (G) Western blot analysis of human EC cell line panel (n=13) with same PAX2 monoclonal antibody used for immunolocalization. Only 2/13 lines (AN3CA and EI) express normal levels of PAX2, consistent with the observed loss in ∼80% of primary EC. (H) *PAX2* mRNA expression levels across human EC lines per qRT-PCR (n*=*3, mean±SEM).

The absence of PAX2 protein in ≥80% of primary EC was consistent with previous findings (*15*). Notably, there was no difference among grade 1 (n=45), 2 (n=37), or 3 (n=33) EC, supporting the idea that PAX2 is an early driver event in EC development (Fig. 1E,F). Western blot analysis of PAX2 of a panel of 13 human EC cell lines revealed that PAX2 protein was undetectable in 9/13 lines and barely detectable in 2/13 (MFE-319 and HEC-1-B) (Fig. 1G). qPCR results were consistent with these findings, suggesting that transcriptional silencing may account for PAX2 protein loss in EC (Fig. 1H). The absence of PAX2 protein in 11/13 (85%) EC lines validated this cell line panel as an experimental system to investigate the origins and functional consequences of PAX2 loss.

### Growth suppression by re-expression in PAX2-deficient EC lines

While loss of PAX2 protein in EIN and EC has been documented in clinical pathology studies, this does not establish a causal link to carcinogenesis. To investigate this, we engineered an inducible lentiviral Tet-On system that permits precise control of *PAX2* expression (Fig. S1A,B). Doxycycline (DOX) treatment resulted in *PAX2* re-expression in PAX2-deficient Ishikawa EC cells (Fig. S1C,D). In vitro growth assays showed that re-expression resulted in a significant suppression of growth (Fig. S1E), and reduced colony formation (Fig. S1F,G). In vivo xenograft assays comparing empty and *PAX2* vector (with DOX-induced expression via drinking water) demonstrated that *PAX2* re-expression resulted in tumor xenografts with slower growth and reduced size (Fig. S1H-J). Western blot analysis and immunolocalization at euthanasia confirmed sustained PAX2 protein expression throughout the experiment (Fig. S1K,L). Similar experiments conducted using another PAX2-deficient EC line, HEC-1-A, yielded comparable results (Fig. S1M-V).

### Growth promotion by *PAX2* knockdown (KD) in a PAX2-expressing line

We investigated the effects of *PAX2* KD using a lentiviral construct in the EC line AN3CA (Fig. S2A-C), which expresses PAX2 (Fig. 1G,H). In contrast to *PAX2* re-expression in Ishikawa cells, *PAX2* KD resulted in increased cell growth (Fig. S2D), and accelerated wound closure in 2D wound assays (Fig. S2E,F). Cell cycle analysis showed that *PAX2* KD affected cell cycle progression, with increased numbers of cells in the S and G2/M phases and a concomitant decrease in cells in the G0/G1 phase (Fig. S2G,H). Xenograft assays comparing scrambled shRNA control with *PAX2* shRNA showed that KD resulted in more rapid tumor growth and larger tumors (Fig. S2I-K). Western blot analysis of tumor lysates at the end of the experiment confirmed stable *PAX2* KD (Fig. S2L). Taken together, these complementary sets of experiments provide preliminary evidence for *PAX2* as a significant endometrial tumor suppressor, whose inactivation promotes EC cell growth.

### Evidence against intragenic rearrangements and for *PAX2* transcriptional silencing as the underlying mechanism for PAX2 inactivation

*PAX2* point mutations are rare in EC (<1% of cases), and cannot explain the high incidence of PAX2 loss (*7*). Furthermore, the majority of rare EC harboring *PAX2* single nucleotide coding variants also exhibit ultramutation due to *POLE* mutations, indicating that these *PAX2* variants are “bystanders” due to high mutational burden (*20*). *PAX3* and *PAX7* chromosomal translocations define childhood rhabdomyosarcomas, raising the possibility that *PAX2* gene rearrangements or deletions might analogously underpin PAX2 expression loss in EC. Break-apart FISH using BAC probes 5’ and 3’ of the gene was conducted in 12 cases of EIN with definitive and complete PAX2 protein loss. In all cases, 5’ and 3’ *PAX2* signals were juxtaposed within the interphase nuclei of EIN epithelial cells, with no loss of either signal. These results ruled out *PAX2* deletions or intrachromosomal rearrangements as the general mechanism accounting for PAX2 inactivation (Fig. 2A).

**Fig. 2.**
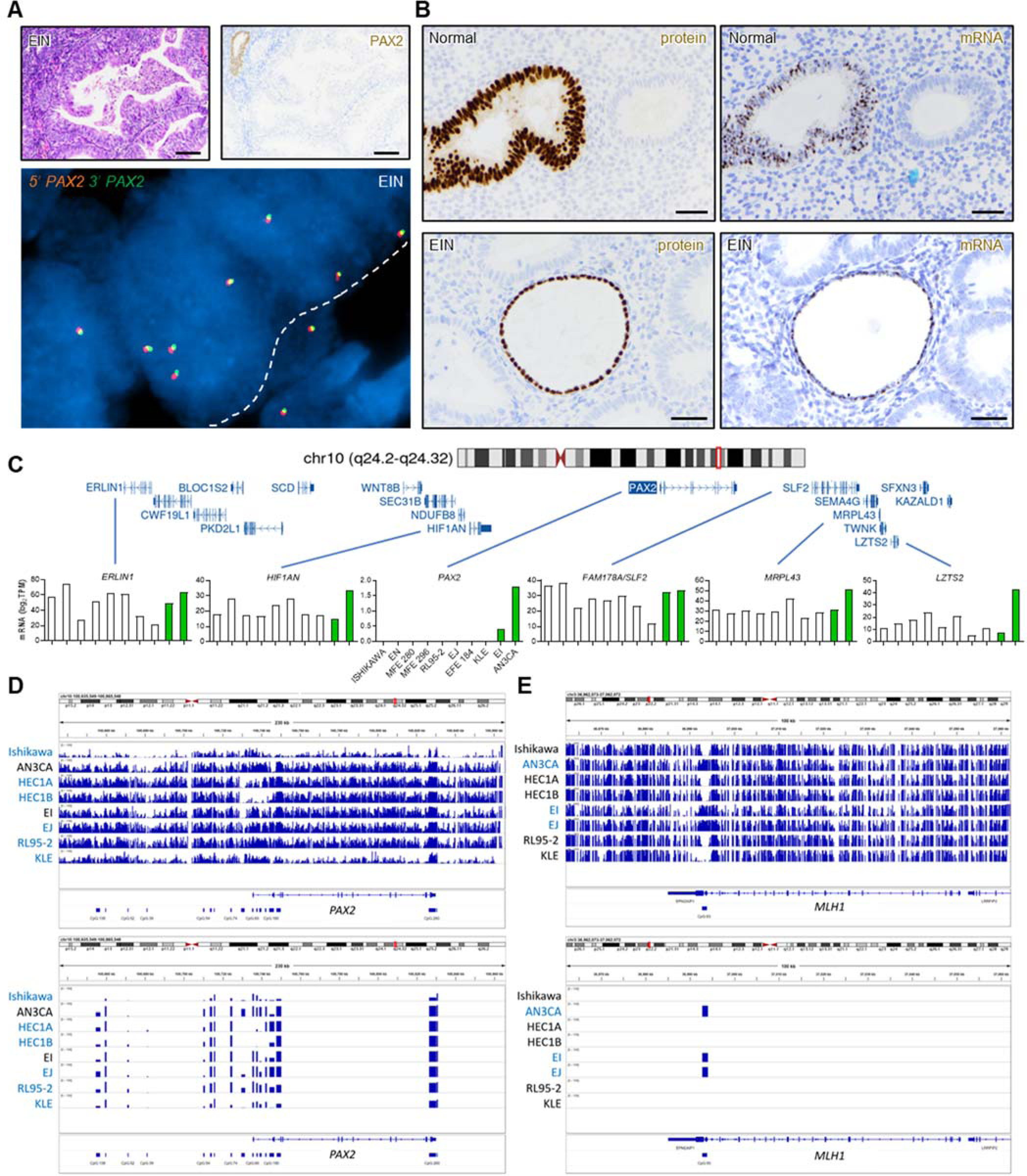
PAX2 protein loss is due to transcriptional silencing specific to *PAX2* locus. (A) Top panels, PAX2-deficient EIN. Single gland of residual normal endometrium serves as internal positive control for PAX2 expression. EIN glands show complete loss of PAX2 protein. Bars=200 μm. Bottom panel, break-apart FISH for *PAX2* locus with flanking BAC probes 162 kbp 5’ (Spectrum Orange) and 188 kbp 3’ (Spectrum Green) from *PAX2* gene body in a PAX2-deficient gland. No absent or physically separate orange and green signals are evident. White dashed line demarcates epithelial/stromal boundary. EIN from n=12 patients analyzed with similar results. (B) Immunolocalization and RNA-ISH of PAX2 loss of expression in serial sections. Top panels, normal human endometrium with single isolated PAX2-deficient gland. Bottom panels, EIN with diffuse PAX2 protein loss. Single entrapped normal (non-neoplastic) gland expresses PAX2 protein (internal positive control). n=6 normal endometria with PAX2-null clones and n=6 EIN with diffuse PAX2 loss were analyzed, with similar results. Bars=50 μm. (C) Expression of individual genes adjacent to *PAX2* locus across EC lines per RNA-seq. Y axis=mRNA abundance as log_2_ transcripts/million (TPM). Both PAX2 expressing EC lines indicated with green bars. (**D-E**) Targeted methyl-seq of *PAX2* (230 kbp) and *MLH1* (100 kbp) coding and flanking genomic regions. CpG islands per UCSC genome browser (GRCh37/hg19) shown for both loci (*47*). Integrated Genomics Viewer (IGV) shows methylation peaks across both loci. Cell lines highlighted in blue are silenced for the respective locus (*PAX2* or *MLH1*). (D) Methyl-seq of *PAX2*. Neither large-nor small-scale methylation events correlated with silencing. (E) Methyl-seq of *MLH1*. Silencing correlated with strong methylation signal in single CpG island known to account for *MLH1* silencing in EC.

Next, to determine whether PAX2 protein loss in endometrium results from transcriptional mechanisms, we performed *PAX2* RNA-ISH (RNAscope) on 6 cases of normal endometria with minute PAX2-deficient clones and 6 cases of EIN. PAX2 immunolocalization was performed on adjacent step sections. Entrapped normal glands served as internal controls. Remarkably, the protein and mRNA loss patterns were superimposable in all 12 cases (Fig. 2B). These findings from human tissue specimens establish that 1) PAX2 protein loss occurs at the transcriptional level (gene silencing), and 2) this gene silencing event represents a very early, if not initiating, driver event in EC genesis.

### *PAX2* silencing is restricted to *PAX2* locus

The *PAX2* gene (∼100 kbp) resides in a ∼350 kbp gene desert. We sought to determine whether *PAX2* silencing occurred across a larger region and whether neighboring genes were transcriptionally perturbed. RNA-seq of 10 EC lines, including two non-silenced lines, showed that *PAX2* was the only silenced locus among its genomic neighbors (Fig. 2C). Additionally, Western blot analysis of *HIF1AN* (5’ neighbor) showed no downregulation in the *PAX2*-silenced lines (Fig. S3A). These and the above results establish that *PAX2* is the sole and specific target of a distinct gene-level epigenetic reprogramming event that initiates most EC.

### Abnormal methylation at *PAX2* locus does not explain PAX2 loss

In EC, hypermethylation of a 5’ *MLH1* CpG island results in locus-specific silencing (*5, 21*). A previous report suggested that the *PAX2* promoter is normally hypermethylated, but becomes unmethylated in EC (*18*), although this is opposite to *MLH1* and other tumor suppressors subject to promoter hypermethylation. Using the same methylation-specific PCR (MS-PCR) assay, we analyzed non-neoplastic endometria from 40 women. *PAX2* was consistently unmethylated; no specimen exhibited predominant methylation of *PAX2* (Fig. S3B,C). Thus, we were unable to reproduce the results of a previous report (*18*). Another study using this MS-PCR assay also found that *PAX2* was unmethylated in normal endometrium (*22*).

MS-PCR evaluates the methylation status of only a few bases within a single CpG island. To overcome this limitation, we performed targeted methyl-seq of 230 kbp encompassing *PAX2* in eight EC lines including the 2 retaining PAX2 expression (Fig. 1G,H). Despite the presence of differentially methylated regions, we did not observe any methylation feature(s) including 1) CpG islands, 2) subregions thereof, or 3) non-CpG regions in the gene body or flanking sequences, which correlated with *PAX2* expression (Fig. 2D). In contrast, methyl-seq of a 100 kb *MLH1* region revealed abnormal methylation at only one 5’ CpG island (Fig. 2E), where hypermethylation was consistently associated with *MLH1* silencing, as confirmed by western blotting (Fig. S3D; see legend for additional detail.). Thus, neither promoter hypermethylation nor abnormal methylation at the *PAX2* locus is the basis of PAX2 inactivation in EC.

### CRISPR-mediated activation establishes an epigenetic basis for *PAX2* silencing and its reversibility

To confirm that another epigenetic mechanism underlies *PAX2* silencing in light of the above unexpected result, we took advantage of CRISPR activation (CRISPRa) using an all-in-one lentiviral vector encoding an endonuclease-deficient mutant Cas9 (dCas9) fused to the transcriptional activator VPR, with a guide RNA (sgRNA) targeting the dCas9-VPR transactivator to *PAX2* (Fig. 3A,B) (*23*). Four sgRNAs targeting *PAX2* were tested, and the guide (#4) with the strongest *PAX2* induction in Ishikawa was chosen (Fig. 3C-E). In all 12 EC lines, transduction of the CRISPRa-*PAX2* lentivirus with puromycin selection increased *PAX2* expression per qPCR. Induction levels varied over several logs, from 4-5x to >1000x relative to non-targeting control lentivirus (Fig. 3F). In all lines, induction of the PAX2 protein was also observed (Fig. 3G). Increased *PAX2* expression was most dramatic in *PAX2*-silenced lines such as Ishikawa and MFE-296. Induction was lower in the non-silenced line EI, as might be expected, but was still significant. Growth assays with Ishikawa and HEC-1-B cells (Fig. 3H) demonstrated that *PAX2* CRISPRa resulted in significant growth suppression comparable to the enforced expression of *PAX2* cDNA (Fig. S1E). These findings further demonstrate that the *PAX2* locus is not irreversibly damaged in EC, as would occur with gene deletions/internal rearrangements. In addition, these results constitute a proof-of-principle that *PAX2* silencing in EC is reversible, with potential therapeutic implications.

**Fig. 3.**
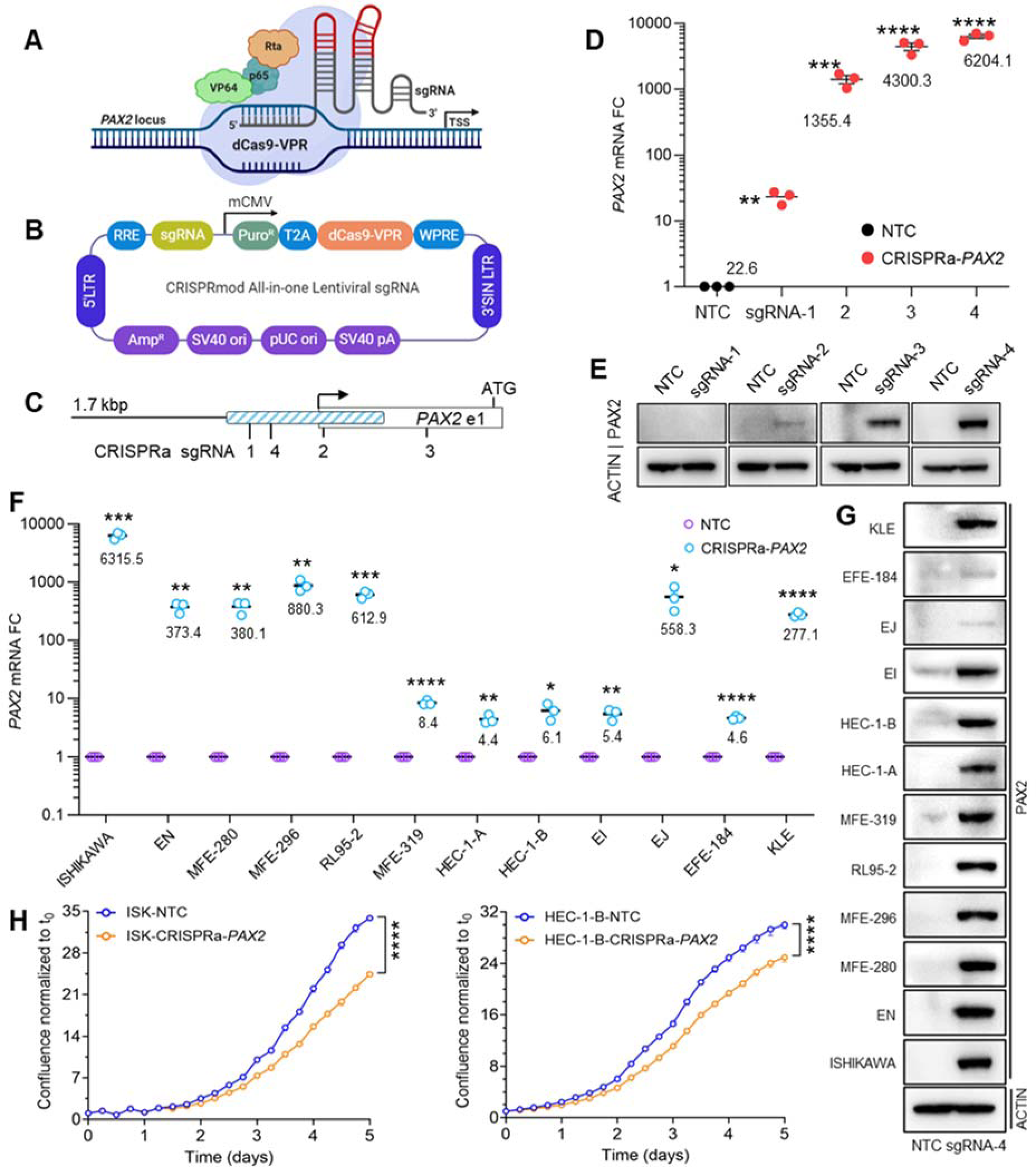
Reversal of *PAX2* silencing by CRISPRa. (A) CRISPRa strategy targeting dCas9-VPR to *PAX2* locus with guide RNA (sgRNA). (B) All-in-one lentiviral construct with sgRNA and dCas9-VPR (created with BioRender.com). (C) Relative positions of four sgRNAs to transcriptional start site (arrow). Start ATG codon in first exon (unfilled rectangle) is shown relative to 1598 bp *PAX2* proximal-promoter enhancer region (blue hatched rectangle). The 20 bp sgRNA#4 is 133 bp 3’ of transcriptional start site (D) *PAX2* mRNA expression by qPCR in EC lines following CRISPRa with non-targeting control (NTC) and *PAX2*-specific sgRNAs (n=3, mean±SEM, multiple t tests). (E) Western blot analysis of PAX2 expression in Ishikawa cells after CRISPRa with four guide RNAs. (F) qPCR of *PAX2* mRNA in Ishikawa cells following CRISPRa with non-targeting control (NTC) and sgRNA #4 (n=3, mean±SEM, multiple t-tests). (G) Western analysis of PAX2 expression in EC lines subjected to sgRNAs #4 CRISPRa. (H) Cell growth following CRISPRa in Ishikawa (ISK) and HEC-1-B by live cell imaging (n=3, mean±SEM, unpaired, 2-tailed t test). For all panels, **P<0.01; *P<0.05; **P<0.01; ***P<0.001; ****P<0.0001.

### *PAX2* silencing is associated with loss of a promoter-proximal active enhancer and gain of facultative heterochromatin features

After eliminating small-scale mutations, genomic rearrangements, and abnormal DNA methylation as causes of *PAX2* silencing in EC lines and primary tumors, we investigated alternative chromatin-based epigenetic mechanisms. We analyzed a 1 Mbp region spanning *PAX2* with published ATAC-seq datasets (GSE114964) for PAX2+/non-silenced (AN3CA) and PAX2–/silenced (Ishikawa, KLE, RL95-2) EC lines (*24*). The most striking difference was in the *PAX2* promoter, where a ∼1.5 kbp region exhibited open chromatin only in the PAX2+ line (Fig. 4A). This led us to hypothesize that this active chromatin feature is unique to PAX2+ cells. To profile enhancer activity in this region, we conducted H3K27ac ChIP-seq on PAX2+ AN3CA vs. PAX2– Ishikawa cells, confirming an active enhancer in AN3CA but not Ishikawa cells (Fig. 4B). This discovery was independently validated using the H3K27ac CUT&Tag assay. Given the overlap of this enhancer with the *PAX2* promoter, we considered this regulatory element to be a promoter-proximal enhancer that governs *PAX2* transcription.

**Fig. 4.**
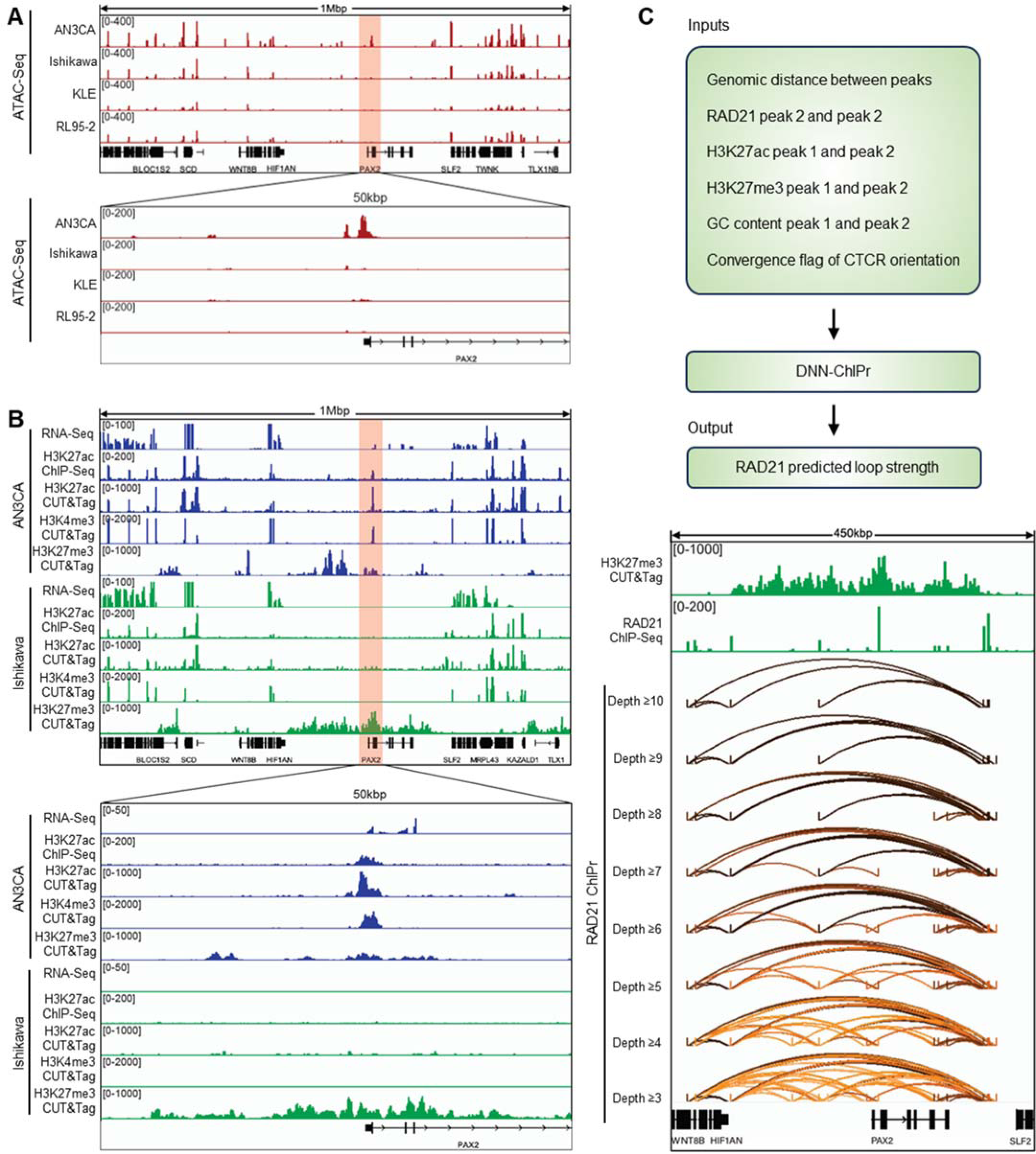
*PAX2* silencing is associated with loss of promoter-proximal active enhancer and gain of facultative heterochromatin features. Proximal promoter enhancer region=chr10:102504680-102506278 (1598 bp) (GRCh37/hg19). **(A)** ATAC-seq analysis of *PAX2* locus in AN3CA, Ishikawa, KLE, and RL95-2 cells. AN3CA cells were PAX2+ (non-silenced), whereas Ishikawa, KLE, and RL95-2 cells were PAX2– (silenced). **(B)** Comprehensive transcriptomic and epigenetic (H3K27ac, H3K4me3, and H3K27me3) profiling of PAX2+ (AN3CA) and PAX2– (Ishikawa) cells. **(C)** Predicted RAD21 ChIA-PET analysis of Ishikawa cells using ChIPr with varying PET interaction strengths, focusing on an H3K27me3 enriched region surrounding *PAX2*. The top panel shows the schematic of the ChIPr pipeline. Interaction strengths are represented by depth values ranging from ≥3 to ≥10.

Consistent with this observation, the *PAX2* promoter was associated with the active promoter mark H3K4me3 in AN3CA but not in Ishikawa cells (Fig. 4B). Furthermore, loss of *PAX2* expression in Ishikawa cells was associated with formation of H3K27me3 domains, representing inaccessible chromatin/facultative heterochromatin; this was markedly less pronounced in PAX2+ AN3CA cells and restricted to regions outside the *PAX2* gene body (Fig. 4B). These results indicate that *PAX2* transcriptional silencing is associated with loss of open/active chromatin marks and gain of inaccessible chromatin/facultative heterochromatin features. Given that *PAX2* transcriptional silencing is the earliest known initiating event in EC, our results provide new insights into the epigenetic basis of this disease.

### Cohesin-mediated 3D genome organization and focal *PAX2* silencing in EC

We further investigated mechanisms underlying *PAX2* silencing and found that it was associated with repressive H3K27me3 marks across the gene desert, but the marks did not spread to neighboring genes, explaining why these *PAX2* neighbors were not transcriptionally affected (Fig. 4B). This indicated that the *PAX2* desert may be insulated from neighboring genes through the formation of an insulated gene neighborhood in the context of the 3D genome (*25, 26*). We hypothesized that the cohesin complex forms an insulated neighborhood via a looping mechanism, restraining the spread of the H3K27me3 domain beyond the desert. To assess this, we analyzed Ishikawa ChIP-seq data for RAD21 (a cohesin complex component) and observed multiple RAD21 peaks in the *PAX2* gene desert (Fig. 4C).

We recently developed Chromatin Interaction Predictor (ChIPr), a machine learning model based on deep neural networks, to predict cohesin-mediated chromatin interaction strengths between any two loci (*27*). Our model uses ChIP-seq signals for RAD21, H3K27ac, and H3K27me3 as inputs and predicts RAD21 chromatin interaction analysis by paired-end tag sequencing (ChIA-PET) as output. We utilized ChIPr to detect the strength of all combinations of RAD21 loops between RAD21 peaks in the *PAX2* desert. Paired-end tags (PETs) are units used for measuring the interaction strength between a pair of anchor peaks, with more PETs between anchors signifying stronger interactions. We used ChIPr to predict all PETs with a depth of ≥3 between the RAD21 peaks (Fig. 4C). Next, we systematically eliminated weak interactions in a stepwise manner by traversing from interactions with PET depths ≥3 to ≥10. This enabled us to identify the strongest cohesin loops and discovered that the *PAX2* desert is insulated from neighboring genes through the formation of a cohesin-mediated insulated gene neighborhood. Remarkably, the spread of the H3K27me3 repressive domain was perfectly contained within the strong cohesin loops, explaining why the outside genes were not silenced (Fig. 4C). Our results indicate that a complex interplay between the 3D genome and the epigenome underlies focal *PAX2* transcriptional silencing in EC models.

We next investigated if the *PAX2* insulated neighborhood is a unique feature of Ishikawa cells or a more generalized feature of all human cells. Analysis of experimental RAD21 ChIA-PET data from 24 human cell types from the ENCODE portal indicated that *PAX2* resides in an insulated neighborhood in all cell lines, making the insulated neighborhood a universal feature of *PAX2* in human cells (Fig. S4) (*28*). Given that the loss of many tumor suppressors (*TP53, PTEN, RB1* etc.) occurs via focal DNA deletions, *PAX2* transcriptional silencing in the context of cohesin loops is a non-genetic/epigenetic equivalent of focal deletions in cancer.

### Mechanisms underlying *PAX2* silencing in EC PDX models

Next, we explored the generalizability of these cell line-based discoveries by analyzing EC patient-derived xenograft (PDX) models. We examined three PDXs: one PAX2+ (PDX441) and two PAX2– (PDX164 and PDX333). We conducted ATAC-seq and examined the same 1 Mbp spanning *PAX2*. Consistent with our cell line models, the most striking difference was observed for the *PAX2* promoter. In the PAX2+ PDX, this region exhibited open chromatin. Unlike PAX2+ AN3CA cells, where the open chromatin was approximately 1.5 kbp, the open chromatin in the PAX2+ PDX spanned >10 kbp (Fig. 5A), indicating characteristic differences between the cell lines and PDX models. Enhancer activity was analyzed by profiling H3K27ac using ChIP-seq and CUT&Tag. Both experiments indicated that the open chromatin identified by ATAC-seq represented a massive enhancer spanning >10 kb, characteristic of a super-enhancer, which was unique to the PAX2+ PDX (Fig. 5A). Consistent with this observation, the *PAX2* super-enhancer exhibited active promoter (H3K4me3) signals in the PAX2+ PDX, but not in PAX2– PDXs. Conversely, PAX2– PDXs exhibited higher enrichment of the repressive H3K27me3 mark in *PAX2*. Taken together, these data indicate that the mechanism of *PAX2* transcriptional silencing is remarkably similar in both cell line and PDX models.

**Fig. 5.**
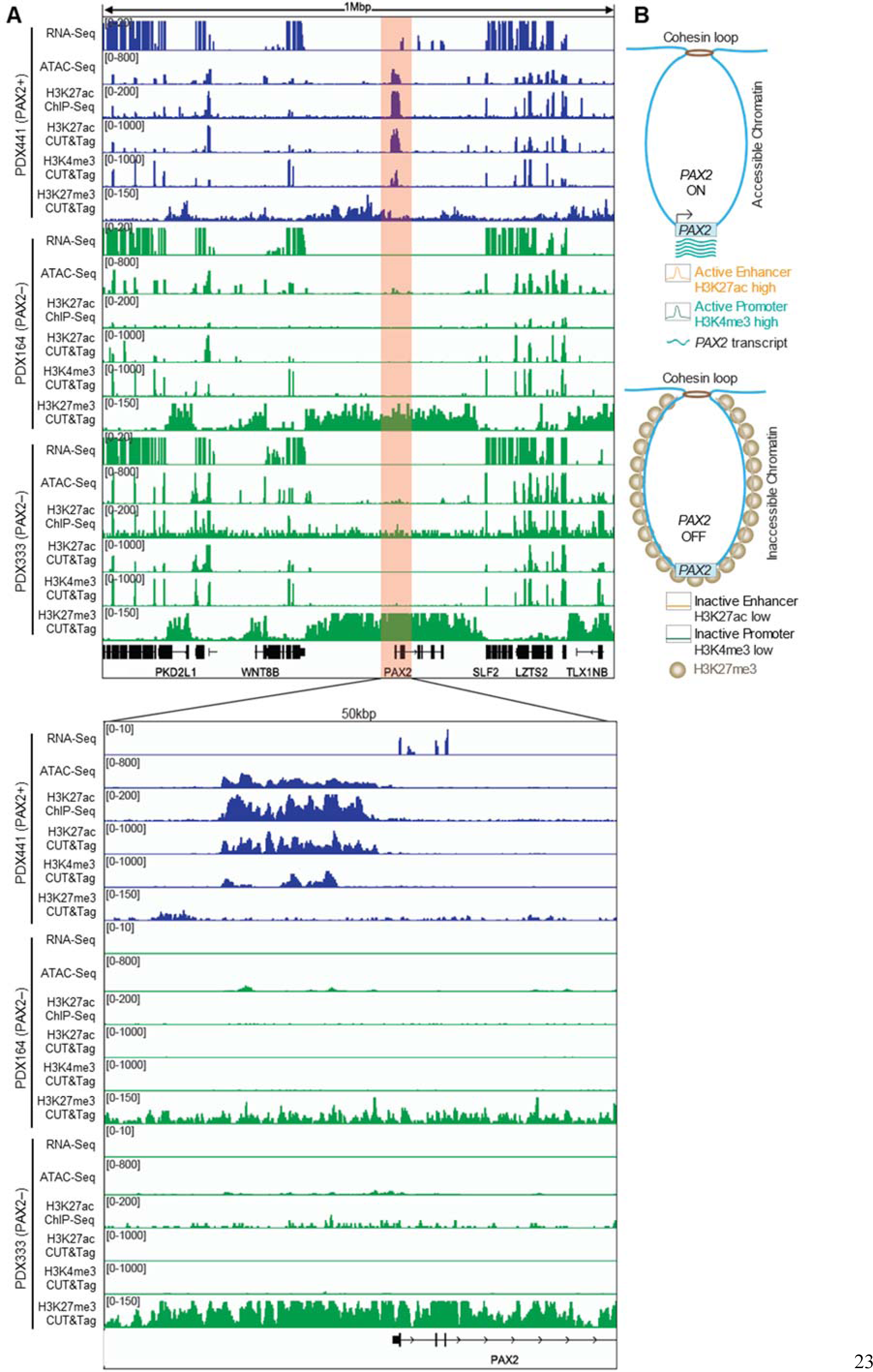
Cohesin-mediated 3D genome organization and focal *PAX2* silencing in EC. (A) Comprehensive transcriptomic and epigenetic (ATAC-seq, H3K27ac, H3K4me3, and H3K27me3) profiling of PAX2+ and PAX2– PDX models of EC. (B) Pearl necklace model of *PAX2* transcriptional silencing.

Based on these observations, we propose a “pearl necklace” model for *PAX2* transcriptional silencing (Fig. 5B). The loss of *PAX2* expression is associated with the replacement of open/active chromatin features (H3K27ac and H3K4me3) with inaccessible chromatin features (H3K27me3). The spread of H3K27me3 signal resembles a pearl necklace, with its length adjusted by cohesins.

### *PAX2* is an oncodevelopmental tumor suppressor regulating endometrial gene expression via control of the enhancer landscape

Prior evidence that *PAX2* is a pioneer TF (*29, 30*) led us to hypothesize that *PAX2* regulates EC transcriptomes by shaping enhancer activity. We conducted H3K27ac ChIP-Seq to compare enhancer profiles in PAX2+ vs. PAX2– (shRNA KD) AN3CA cells and PAX2– vs. PAX2+ (re-expressed) Ishikawa cells. In AN3CA, we identified ∼17.5K H3K27ac peaks with *PAX2* downregulation resulting in both gain and loss of thousands of enhancers (Fig. 6A,C). Similarly, re-expression of *PAX2* in Ishikawa cells also resulted in both gain and loss of thousands of enhancers (Fig. 6B,D). These results indicated that changes in *PAX2* status shaped the chromatin landscape by commissioning and decommissioning enhancers. Notably, these enhancer alterations were predominantly in distal regulatory regions, rather than gene promoters (Fig. 6A,B). We hypothesized that changes in the activity of these distal enhancers contribute to transcriptomic dysregulation via long-range chromatin interactions.

**Fig. 6.**
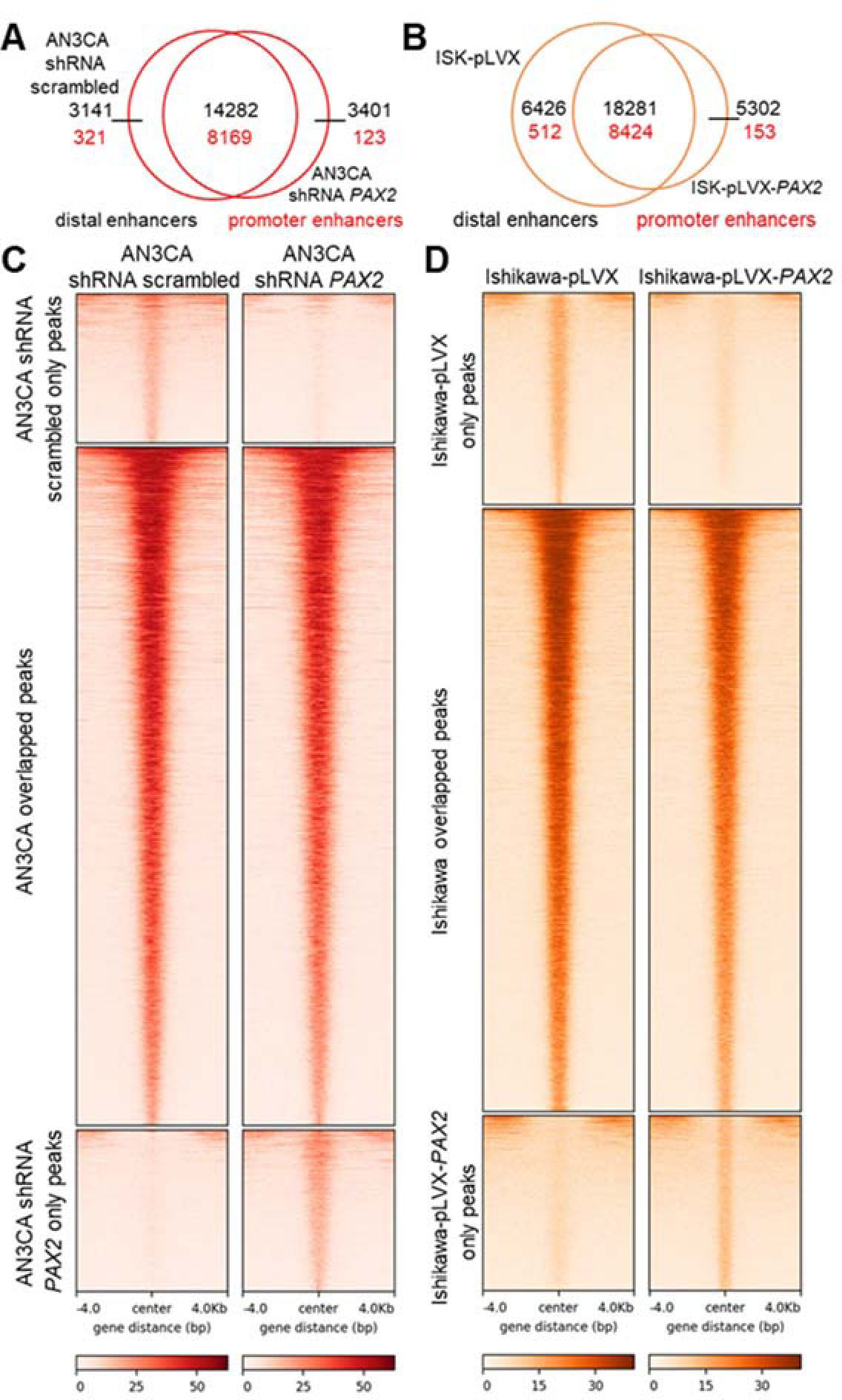
*PAX2* knockdown (KD) and re-expression alters enhancer profiles per H3K27ac ChIP-seq. (**A-B**) Venn diagrams of H3K27ac ChIP-seq; enhancer peaks in AN3CA **(A)** (scrambled and *PAX2* KD) and Ishikawa cells **(B)** (control and *PAX2* re-expressed) in the promoter and distal regions. (C) In AN3CA cells, scrambled shRNA peaks were exclusively identified in scrambled shRNA-treated cells. Overlapping peaks were common between scrambled and *PAX2* KD cells, and shRNA *PAX2* peaks were found in *PAX2* KD cells. (D) In Ishikawa cells, empty vector (pLVX) peaks were exclusively present in pLVX-treated cells, overlapping peaks were common between pLVX- and *PAX2*-expressing cells (pLVX-*PAX2)*, and *PAX2* re-expression peaks were identified in pLVX-*PAX2* cells.

Re-expression of *PAX2* in Ishikawa cells resulted in hundreds of differentially expressed genes (DEGs) (246 upregulated and 352 downregulated genes, p,q<0.05, -1<log_2_FC>1), indicating transcriptional reprogramming. Gene ontology analyses revealed statistically significant enrichment of genes involved in anatomical structure development, developmental processes, and tube development, underscoring PAX2’s role as an oncodevelopmental factor with broad functional impacts (Fig. S5A). Although PAX2’s expansive impacts on the enhancer landscape and transcriptional reprogramming rationalize its activity as an endometrial tumor suppressor and argue against the overriding significance of individual genes, *PGR* stood out as a potentially significant target, whose expression significantly increased (6.98x, p,q <0.00001) after *PAX2* re-expression in Ishikawa (Fig. S5B). In ISK-pLVX-*PAX2* cells, progesterone receptors encoded by *PGR* (PR-A/B) were increased at the protein level (2.2x) (Fig. S5B,C). To further validate the functional impact of increased PR-A/B, we treated Ishikawa cells (PAX2+/–) with progesterone and observed modest, albeit significant, suppression of cell growth (Fig. S5D). These findings are consistent with prior results linking PAX2 to the transcriptional regulation of *PGR* (*31*) and are further explored in mouse models below.

### Mouse model establishes *Pax2* as in vivo EC tumor suppressor that synergizes with *Pten*

While the above studies provided evidence for a tumor suppressor role of *PAX2* in EC, cell lines have limitations as experimental model systems, including genetic divergence from original tumors and absence of tumor microenvironment plus other host factors (*32*). To overcome these limitations and test the hypothesis that *PAX2* is an EC tumor suppressor in the most rigorous in vivo genetic system, we explored the biological functions of *Pax2* in genetically engineered mice.

We utilized endometrial epithelium-specific *BAC-Sprr2f-Cre*, which becomes active after sexual maturity at 5 weeks of age (*33, 34*), and floxed (L) *Pax2* and *Pten* alleles. *Pax2^L/L^* mice are viable, whereas *Pax2^del/del^* embryos exhibit renal agenesis, confirming that *Pax2^L^* yields a null mutation following Cre-mediated gene ablation (*35*). *BAC-Sprr2f-Cre; Pax2^L/L^*(*Pax2*), *BAC-Sprr2f-Cre; Pten^L/L^* (*Pten*), and *BAC-Sprr2f-Cre; Pax2^L/L^; Pten^L/L^* (*Pax2/Pten*) females were generated by breeding. *Pten* was selected because 1) single tumor suppressors usually do not yield overt ECs (*33, 34, 36*), 2) *PTEN* is the most frequently mutated gene in EC, and 3) *PAX2* silencing and *PTEN* mutations frequently co-occur in EIN and EC (*14*).

Consistent with prior studies, endometrial-specific *Pten* inactivation resulted in EIN, with only some *Pten* mice developing lethal adenocarcinomas with very long latency (*34, 36-38*). In contrast, *Pax2/Pten* females exhibited a striking and lethal phenotype with early mortality due to EC (Fig. 7A,B). Tumors were classified into two discrete histotypes: endometrioid (16/30 mice, 53.3%), endometrioid mucinous with squamous differentiation (confirmed by p63; 1/30, 3.3%), or an admixture of both (13/30, 43.3%) (Fig. 7C,D). Mucinous and/or squamous differentiation are common features of human EC. Thus, while two distinct tumor histotypes were observed in *Pax2/Pten* EC, often together, both fell within the spectrum of human EC. Immunolocalization confirmed loss of PTEN and PAX2 in malignant epithelial cells (Fig. 7E). *Pax2/Pten* ECs were highly invasive, often into the abdominal cavity, with frequent metastases in 30 mice subjected to complete necropsy, including ovary (60%), kidney (16.7%), liver (10%), pancreas (16.7%), spleen (10%), and intestine (33.3%), with distant metastases in lymph nodes (6.7%) and lung (16.7%) (Fig. 7F).

**Fig. 7.**
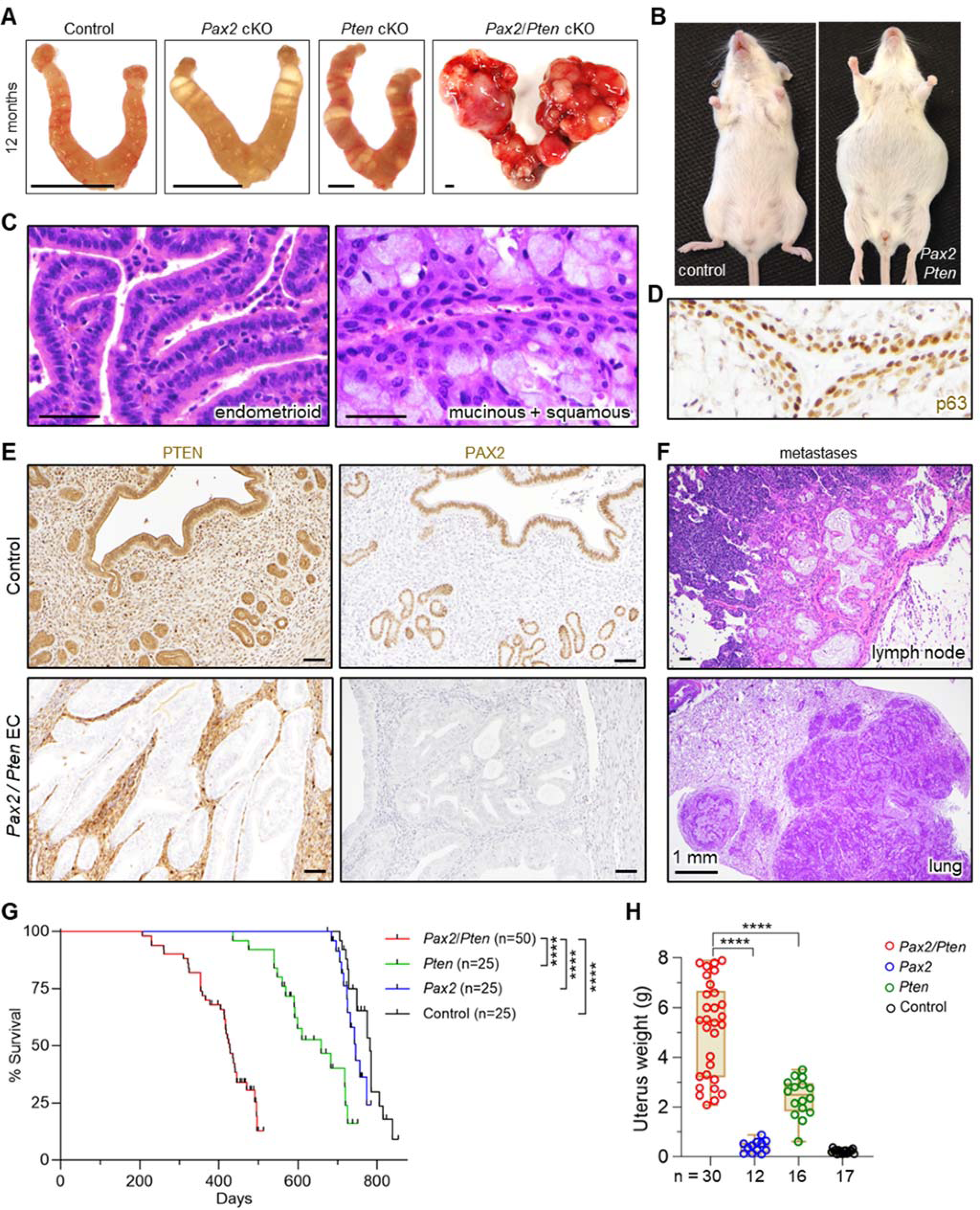
*Pax2* is EC tumor suppressor synergizing with *Pten* in vivo. **(A)** Uteri at 12 months of age. Bars=2 mm. **(B)** *Pax2/Pten* mouse with distended abdomen due to tumorous uterus and ascites. This phenotype was not observed in single knockouts. **(C)** Distinct EC histotypes in *Pax2/Pten* females, H&E. Bars=50 μm. **(D)** p63 immunostaining in mucinous+squamous EC confirming squamous differentiation. **(E)** PTEN and PAX2 immunostains confirming endometrial-specific ablation in invasive *Pax2/Pten* EC. Bars=50 μm. **(F)** Distant metastases from *Pax2/Pten* EC, H&E Bar=50 μm. **(G)** Survival analysis of *Pax2*/*Pten* (n*=*50), *Pten* (n*=*25), and *Pax2* (n*=*25), and littermate control (n*=*25) mice; ****P<0.0001 per log-rank test. **(H)** Uterine weights at necropsy, x axis shows number of animals per genotype. ****P<0.0001, 1-way ANOVA, Tukey’s multiple-comparison test. Bars=50 μm.

*Pax2* mice showed a minor decrease in survival, with only a subset displaying early signs of invasive EC; however, this was not statistically significant. In contrast, *Pax2*/*Pten* mice had significantly shorter median survival than littermate controls or single knockout mice (Fig. 7G). Uterine weights also confirmed striking cooperativity. While *Pax2* mice had normal uterine weights and *Pten* mice had increased uterine weights due to longer uterine horns rather than invasive cancers (*34*), *Pax2*/*Pten* mice had far higher uterine weights reflecting overt tumor burden (Fig. 7H). In summary, these results provide formal genetic evidence that *Pax2* is an in vivo EC tumor suppressor that synergizes with *Pten,* establishing mice as a useful model for additional investigations into the biology of *Pax2* +/– *Pten* in EC.

### scRNA-seq reveals PAX2-null population and validates *PGR* as *PAX2* target

Our EC analyses indicated that *PAX2* regulates *PGR*. Previous studies have shown that among established EC lines, only Ishikawa express ERα and PR-A/B (*36, 39*). To explore whether loss of *PAX2* is associated with a reduction in *PGR* in vivo, we performed single-cell RNA sequencing (scRNA-seq) of *Pax2*-mosaic uteri at 8 weeks (see next section and Fig. S7A for explanation of mosaic system). UMAP plots revealed diverse uterine populations, including stromal cells, together with glandular and luminal epithelial cells, among other cell types (Fig. 8A). Violin plots for selected informative genes are shown in Fig. 8B; for example, *Foxa2* distinguishes luminal from glandular epithelial cells (*36*), whereas *Krt8* marks both luminal and glandular epithelial cells. The identification of a distinctive *Pax2*-null epithelial cell cluster permitted differentially expressed gene (DEG) analyses relative to PAX2*+* luminal and glandular epithelium, both of which identified *Pgr1* as underexpressed in the PAX2– epithelial population (Log_2_FC -1.74, P=0.02 for glandular epithelium), consistent with the EC line data implicating *Pgr* as one of many *Pax2* targets (Fig. 8C-F). Immunofluorescence of tissue sections from mosaic uteri showed that ERα and PR-A/B were underexpressed in PAX2– cells relative to their PAX2+ neighbors, whereas controls had uniform expression levels of both factors, confirming that *Pax2* regulates their expression (Fig. 8G,H).

**Fig. 8.**
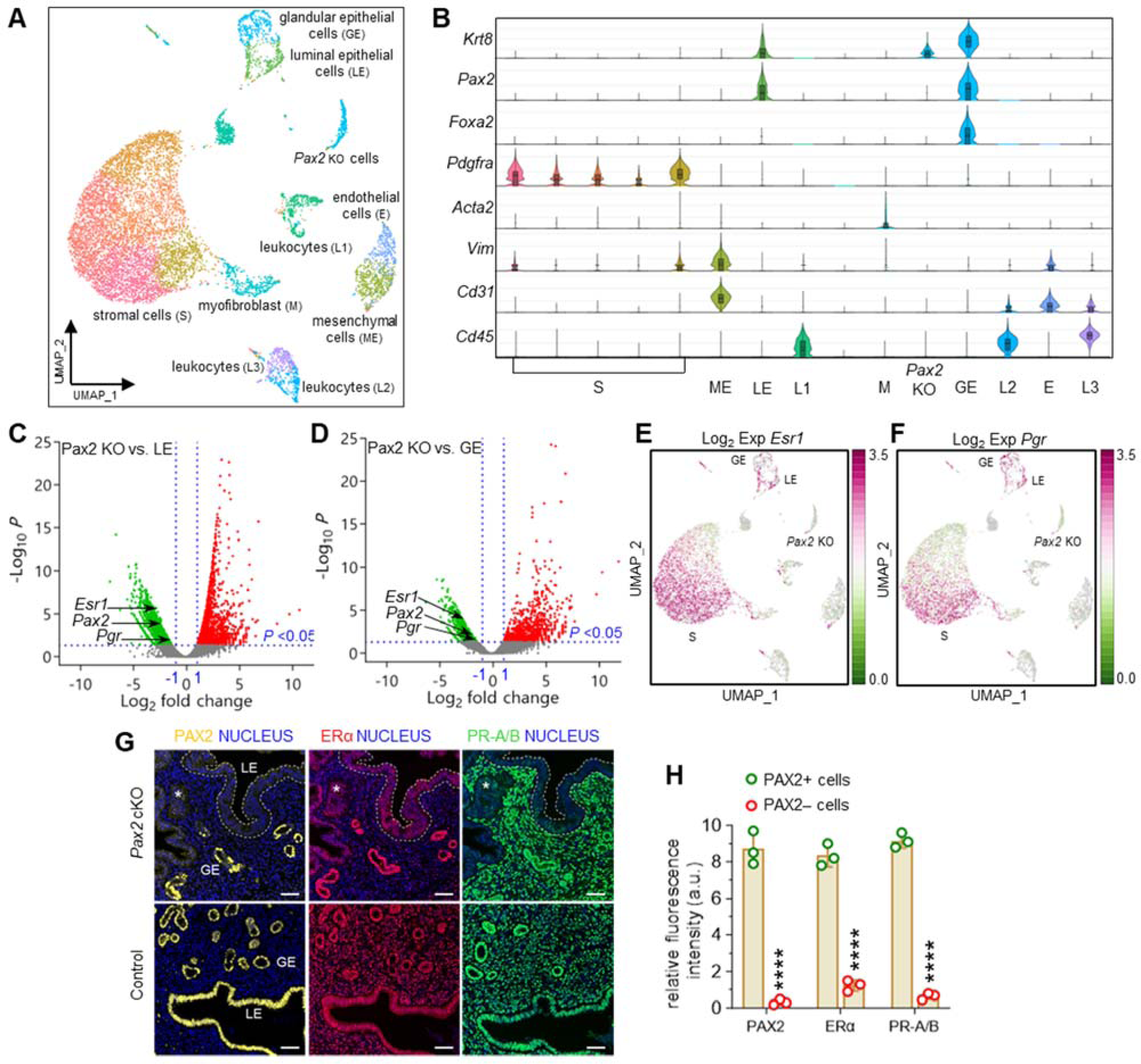
scRNA-seq reveals that inactivation of *Pax2* loss correlates with reduction of *Esr1* and *Pgr* expression in mouse endometrium. Studies were performed with *Pax2*-mosaic uteri at 8 weeks of age. **(A)** Uniform Manifold Approximation and Projection (UMAP) visualization showing cells from *Pax2* mouse uterus (n=2) clustered into 15 distinct subpopulations based on established lineage markers. **(B)** Stacked violin plots show expression of gene signatures associated with known uterine cell types, facilitating identification of lineages within clusters. **(C-D**) Volcano plot showing DEGs (P<0.05) in *Pax2* KO cluster compared to both luminal epithelial (C) and glandular epithelial cell clusters (D). Vertical dotted lines represent log_2_ fold change threshold of ±1, while the horizontal dotted line represents a P-value threshold of 0.05. Selected genes shown. **(E-F)** UMAP plots of *Esr1* (E) and *Pgr* (F). **(H)** Immunofluorescence staining for PAX2, Erα (*Esr1*), and PR-A/B (*Pgr*) in control and *Pax2* mouse uterine adjacent sections (n=3). GE: glandular epithelium; LE: luminal epithelium. Bars=50 μm. **(I)** Comparison of relative fluorescence intensities of PAX2, ERα, and PR-A/B between PAX2+ and PAX2– cells. Data are shown as mean±SEM (n=3); ****P<0.0001, multiple 2-tailed t tests.

### Organoids reveal novel, synergistic growth phenotypes in 3D culture

Epithelial organoids were isolated from control, *Pax2*, *Pten*, and *Pax2/Pten* mice at 12-16 weeks, before the onset of malignancy at 30 weeks. At this time point, organoids should reflect phenotypes associated with specific engineered mutation(s) rather than the acquisition of additional mutations. Remarkably, *Pax2* inactivation alone produced a distinct phenotype of larger organoids with intraluminal growth, resulting in solid organoids versus controls, which formed single-layered, hollow structures (Fig. S6A,B). This was more clearly observed in serial sections of organoids obtained by confocal z-stack imaging (Fig. S6C). *Pten* inactivation resulted in even larger organoids with hollow lumina, as described previously (*34, 36*). In contrast, *Pax2*/*Pten* organoids were significantly larger than the single knockouts (Fig. S6A-C). Moreover, these double-knockout organoids also exhibited aberrant growth into lumina, suggestive of EMT phenotypes (*34*). These synergistic growth patterns were evident as accelerated growth and increased cell numbers in 3D culture (Fig. S6D,E).

The *Pax2* single-knockout organoid phenotype is remarkable, as it shows that Pax2 loss alone confers cellular phenotypes that favor growth. To further explore this, we took advantage of mosaic patterns of *Pax2* ablation resulting from subtotal *BAC-Sprr2f-Cre*-mediated recombination in young females, leading to coexistence of mutant PAX2– and PAX2*+* cells within glands (Fig. S7A,B). Initially, organoids from *Pax2* mice at 8 weeks of age contained substantial proportions of PAX2+ cells (∼35-40%) (Fig. S7C,D). However, after serial passaging, PAX2+ cells rapidly declined and disappeared by 3^rd^ passage (Fig. S7C,D). Since the culture medium lacked estradiol (required for *BAC-Sprr2f-Cre* expression), ex vivo loss of PAX2 expression was not likely due to sustained Cre activity (*34*). Control organoids exhibited 100% PAX2 expression throughout serial passages (Fig. S7C,D), as confirmed by confocal z-stack imaging (Fig. S7G). The loss of PAX2+ cells was further validated using *Pax2* qPCR (Fig. S7E,F). These results demonstrated a significant competitive growth advantage of PAX2 over PAX2*+* cells, further rationalizing the emergence of PAX2-null clones in human endometrium.

Taken together, these studies highlight two key findings: 1) *Pax2* regulates endometrial cell growth in vivo, and 2) potent synergism between *Pax2* and *Pten* significantly affects growth and tumor phenotypes. These observations also rationalize the observed loss of PAX2 in EC and its frequent co-occurrence with *PTEN* mutations/PTEN protein loss.

## DISCUSSION

Cancer is driven by cell-heritable alterations that promote abnormal growth and insensitivity to physiological growth control mechanisms. Most documented cancer-driving events are DNA-level mutations, reflecting the ease with which such mutations have been reliably identified at the genomic level through DNA sequencing. One insight from these studies has been the identification of recurring oncogenic mutations in genes controlling chromatin architecture and gene expression, establishing deregulation of epigenetic control mechanisms as a hallmark of cancer (*11, 40*). Mutations in chromatin regulatory factors have broad and pleiotropic effects that alter transcription at the genome-wide level, a phenomenon that should be distinguished from non-mutational epigenetic reprogramming events targeting single loci.

DNA methylation at CpG dinucleotides was the first epigenetic mark identified. CpG dinucleotides are abundant near transcriptional start sites of housekeeping genes, and such promoter “CpG islands” are almost always unmethylated. While CpG methylation is considered an epigenetic mark, it does involve chemical modification of DNA, which distinguishes it from other types of epigenetic alterations based on histone codes. Many tumor suppressor loci, especially those that are also broadly expressed housekeeping genes, harbor CpG promoter islands (*41*). Through yet unknown mechanisms, promoter CpG islands of some tumor suppressor loci become hypermethylated, leading to transcriptional gene silencing through recruitment of repressor proteins, chromatin remodeling, or blocking transcription factor binding. Such hypermethylation events appear to occur in a single cell that then gains a clonal growth advantage, with promoter hypermethylation status stably maintained by DNA methyltransferases during cell division and tumor growth (*11*).

PAX2 is expressed in only a small number of tissues and cell types, including the parathyroid and genitourinary tract (kidney, seminal vesicle, and uterus), where it serves critical functions in organogenesis and development. Unlike ECs, renal cell carcinomas retain PAX2 expression (*42*). PAX8 is also highly expressed in endometrium but does not undergo loss in EC (*43*), making PAX2 silencing a distinctive signature lesion of the endometrium and, as far as is known, unique among the PAX family as a tumor suppressor in the female reproductive tract or elsewhere. PAX2’s status as a tissue-specific developmental factor is consistent with our results that promoter hypermethylation is not the underlying mechanism for PAX2 silencing, given that most tumor suppressor loci subject to CpG island hypermethylation are broadly expressed housekeeping genes.

*PAX2* transcriptional silencing in EC is comparable to *ERG* transcriptional upregulation in prostatic adenocarcinoma, which is strikingly similar to EC. Aging is the primary risk factor for EC and prostate cancer. Both exhibit precancerous histological counterparts, EIN and prostatic intraepithelial neoplasia (PIN), driven by transcriptomic dysregulation. Upregulation of ETS TFs (primarily ERG) and downregulation of *PAX2* are observed in ∼20% of PIN and ∼80% of EIN lesions, respectively, and facilitate transition to carcinoma (*44*). In the prostate, upregulation of ETS TFs and PI3K pathway activation cooperatively drive the transition from PIN to prostate adenocarcinoma (*45, 46*). Likewise, in the context of the endometrium, we now show in our mouse models that inactivation of *Pax2* together with PI3K pathway activation (via *Pten*, the most frequently mutated gene in EC) cooperatively drives the transition from EIN to EC.

Our study shows that a combination of 1) loss of open/active chromatin marks and 2) gain of inaccessible chromatin/facultative heterochromatin features in a framework dictated by cohesin-mediated 3D genomic architecture underlies focal *PAX2* silencing. Although developed to explain *PAX2* transcriptional silencing, we speculate that our pearl necklace model could be generalized to other cancer drivers. Our discoveries open new questions for the field–what are the upstream triggers for the loss and gain of mutually exclusive H3K27ac and H3K27me3 signals, respectively? Given the intricate association of the endometrium with temporal (long- and short-term) changes in hormonal signaling, we speculate that depletion of master transcription factors in the *PAX2* enhancer may lead to H3K27ac signal loss. This may be stochastic or linked to a normal aging process. We also note that when *PAX2* was expressed, the locus was partially bivalent in terms of H3K27ac and H3K27me3 signals. In particular, H3K27me3 signal was not completely lost (Fig. 4B and 5A). Therefore, transient loss of H3K27ac signals can result in the gain of H3K27me3 signals. However, as H3K27me3 contributes to compact and inaccessible chromatin, reestablishment of the H3K27ac signal may become less likely from a biochemical standpoint. This epigenetic switch from H3K27ac to H3K27me3 is likely to undergo positive selection as we have shown that *PAX2* is a tumor suppressor. The cohesin loops serve as guard rails to prevent this biochemical event from spilling over to neighboring genes. Another question is how a stochastic chromatin remodeling event silences *PAX2* in a single cell, which then expands into a minute PAX2-deficient clone. Silencing one copy of *PAX2* may provide a small growth advantage. If so, one *PAX2* copy may be silenced initially, followed by a second stochastic event involving the other allele to completely silence *PAX2*. Alternatively, both alleles may be silenced simultaneously via unknown mechanisms.

Not surprisingly, it has proven difficult to pharmacologically reconstitute the activity of missing or inactive tumor suppressor proteins, making classical tumor suppressors such as TP53 and PTEN ineffective targets. Nonetheless, observations from this study support PAX2 as an actionable target. First, PAX2 loss defines the large majority of primary and metastatic EC, making PAX2-based strategies have a broad potential clinical impact. Second, although heterogeneity is a major factor limiting treatment efficacy, PAX2 loss is an initiating event and usually a molecular feature across ECs. Third, our CRISPRa studies showed that PAX2 was reactivable in all EC lines, and this had phenotypic consequences, confirming that the locus was not irreversibly damaged in EC. Fourth, the CRISPRa results represent a proof-of-principle that PAX2 could be reactivated pharmacologically. Small molecule inhibitors of diverse epigenetic modifier enzymes may lead to the reactivation of PAX2 (*10*), and novel agents could be identified through systematic chemical screening for *PAX2* re-expression (*5*).

In summary, this study established a specific *PAX2* epigenetic reprogramming event as a novel, and highly recurring cancer-initiating mechanism in EC. We have developed a number of resources, including cell lines, PDXs, epigenomic datasets, and a genetically engineered mouse model that we employed to answer fundamental questions, but are well suited for future investigations to explore further details about *PAX2*’s function as a tumor suppressor or its interactions with PI3K/PTEN and other cancer-causing pathways. These findings and novel approaches have diverse implications for the diagnosis and clinical management of EC, a common, but underestimated malignancy in women.

## Data availability

The data supporting the findings of this study, including Methyl-Seq, RNA-Seq, scRNA-Seq, ATAC-Seq, ChIP-Seq, and CUT&Tag-Seq datasets, were deposited in the NCBI’s Gene Expression Omnibus (GEO) database and are accessible through accession numbers GSE275208, GSE275345, GSE275320, GSE275221, GSE275222, and GSE275223.

## Acknowledgments

The authors thank Dr. Cheryl Lewis and UT Southwestern Tissue Resource, a shared resource of the Simmons Comprehensive Cancer Center, supported in part by National Cancer Institute award 5P30CA142543. We also thank Scott A. Tomlins for helpful comments on the manuscript.

## Author contributions

Conceptualization: DHC, RSM, SSS, SGR

Data Curation, Formal Analysis, and Visualization: SSS, SGR, YG, SL, AA, XZ, AK, CX

Funding Acquisition: DHC, RSM

Investigation and Methodology: SSS, SGR, ICC, PK, SR

Project Administration and Supervision: DHC, RSM, CX

Resources: SR, RRB, VLB-J, ABG, JL, EL, AK

Writing Original Draft, Review and Editing: DHC, RSM, SSS, SGR

