## Supplemental materials for "PAX2 is Transcriptionally Silenced by a Distinct Mechanism of Epigenetic Reprogramming to Initiate Endometrial Carcinogenesis"

Running title: PAX2 transcriptional silencing in endometrial cancer

The authors declare no potential conflicts of interest.

Corresponding authors: Diego H. Castrillon and Ram S. Mani, Department of Pathology, UT Southwestern, 6000 Harry Hines Boulevard, Dallas, TX 75390-9072.

### MATERIALS AND METHODS

#### Nomenclature

Per standard HUGO nomenclature, in this manuscript, *Pax2*=mouse gene or mRNA, *PAX2*=human gene or mRNA, PAX2=mouse or human protein.

#### Human tissues

For *PAX2* methylation PCR analysis, immunohistochemistry, and in situ hybridization, human endometrial tissue sections (both normal and cancerous) were obtained from formalin-fixed paraffin-embedded tissue blocks (UT Southwestern Clinical Pathology Laboratories) using an IRB-approved protocol. Normal archival formalin-fixed paraffin-embedded (FFPE) endometrial specimens for age studies were retrospectively identified from biopsies for workup of abnormal uterine bleeding and where the histologic diagnosis was proliferative endometrium (the preovulatory phase of the menstrual cycle when the endometrium is mitotically active). Abnormal endometrial architecture or history of endometrial neoplasia were exclusion criteria. Tissue microarray sections (US Biomax) from endometrial cancer (EC) patient samples were utilized for PAX2 immunohistochemistry and determination of H-scores in grade 1-3 EC.

#### Mouse strains and survival analysis

All animal procedures and experiments were conducted in accordance with the guidelines and regulations approved by the UT Southwestern Medical Center Institutional Animal Care and Use Committee. Endometrial epithelium-specific *Pax2* and/or *Pten* homozygous conditional knockout mice were generated by breeding mice harboring floxed *Pax2* (maintained on a 129/Sv × C57BL/6J mixed background) (*1*) and *Pten* (*Pten*$^{tm1Hwu}$/J, stock #004597) (*2*) alleles with *BAC-Sprr2f-Cre* mice (maintained on an FVB background, but now available in a pure C57/B6 background through the Jackson Laboratory as [B6(FVB)-Tg (*Sprr2f-cre*)2DCas/J, Stock #037052] (*3*). Sibling progeny inheriting either a single floxed *Pax2* or *Pten* allele but lacking the *BAC-Sprr2f-Cre* transgene were used as controls. The mice were housed in a pathogen-free animal facility in individually ventilated cages and fed ad libitum on a standard chow diet under a 12:12h light-dark cycle. Survival analyses were conducted on both experimental and sibling control mice selected at the time of weaning.

#### Cell lines

Human EC cell lines Ishikawa (Sigma #99040201), HEC-1-A (ATCC #HTB-112), HEC-1-B (ATCC #HTB-113), AN3CA (ATCC #HTB-111), RL95-2 (ATCC #CRL-1671), KLE (ATCC #CRL-1622), EN (DSMZ #ACC 564), MFE-319 (DSMZ #ACC 423), MFE-280 (DSMZ #ACC 410), MFE-296 (DSMZ #ACC 419), EFE-184 (DSMZ #ACC 230), EI and EJ (obtained from M. Takayama) (*4, 5*) were cultured in MEM/RPMI/DMEM-F12 medium (Gibco) per ATCC recommendations supplemented with 10% heat-inactivated fetal bovine serum (FBS; Sigma) and antibiotics (50 U/mL penicillin and 50 mg/L streptomycin; Thermo Fisher Scientific) at 37°C in a humidified atmosphere with 5% $CO_2$. The cell lines were routinely tested to confirm their *Mycoplasma*-negative status using a Mycosensor PCR assay kit (Agilent Genomics #302108).

#### Lentiviral transduction including enforced *PAX2* expression, shRNA, and CRISPRa

To induce *PAX2* expression in human EC cell lines, *PAX2* cDNA was subcloned from the pCMV6-Myc-DDK vector (OriGene) into the pLVX-Tight vector (Clontech), which is a tetracycline (Tet)-

inducible lentivector. Briefly, *PAX2* cDNA was amplified from a commercial human cDNA clone in pCMV6 (OriGene #RC212051, Accession: NM_003987, transcript variant a) using sequence-specific primers for *PAX2* (Forward: 5'-GGGAATTCTTAAACCTTATCGTCGTC-3' and Reverse: 5'-GGGAATTCTTAAACCTTATCGTCGTC-3'). The amplified *PAX2* cDNA was purified, digested with *EcoRI* (New England Biolabs), and cloned into the pLVX-tight vector. Sanger sequencing of the resulting pLVX-Tight-*PAX2* plasmid confirmed the expected sequence for the *PAX2* cDNA insert. The pLVX-Tight vector (puromycin-resistant), either empty or containing the subcloned *PAX2* cDNA, was transfected into 293T cells to produce the lentivirus. These lentiviral particles were used to infect Ishikawa and HEC-1-A cells. For doxycycline-induced *PAX2* expression, Ishikawa and HEC-1-A cells were infected with lentivirus generated by transfection of the pLVX-Tet ON vector (Clontech, neomycin resistant) into 293T cells. Stable cell lines were established using puromycin (1 μg/mL; Thermo Fisher Scientific) and geneticin (200 μg/mL; Gibco) selection. The cells were cultured in MEM (for Ishikawa) and McCoy's 5A (for HEC-1-A) medium supplemented with 10% Tet System Approved FBS (TaKaRa) and then treated with doxycycline (25 ng/mL; TaKaRa) to induce *PAX2* expression. *PAX2* was also re-expressed in human EC cell lines by employing CRISPRmod All-in-one Lentiviral sgRNA with a dCas9 transcriptional modulation vector (CRISPRa-dCas9; Horizon Discovery #VSGH12558-EG5076), following the manufacturer's protocol. To knockdown *PAX2*, AN3CA cells were transduced with either an empty vector, pGFP-C-shLenti (OriGene #TR30021V), or a vector containing a 29-mer *PAX2*-specific shRNA (GTCAAGTCGAGTCTATCTGCATCCACCAA, OriGene #TL310599V), followed by puromycin selection.

**Development of EC Patient-Derived Xenograft (PDX) tumors**

Primary tumor samples from EC patients were obtained during hysterectomy at the UT Southwestern Medical Center or the University of North Carolina School of Medicine. The samples were collected with informed consent from all the patients and were approved by the Institutional Review Board. Patient tumor tissues were washed in sterile PBS and aseptically cut into 2 × 2 mm pieces (*6*). Female NOD/scid/gamma mice (Jackson Laboratory), aged 6-8 weeks, were anesthetized and a 0.5 cm incision was made near their flanks. Four to five tumor pieces mixed with Matrigel were subcutaneously implanted into the incision pocket. Tumor growth was monitored weekly, and once the tumors reached approximately 10 mm in diameter, the mice were euthanized and the tumors were harvested. Chromatin and RNA were extracted from the tumors for ATAC-Seq and RNA-Seq analyses. Additionally, the tumors were processed for FFPE and PAX2 immunohistochemistry to assess the presence or absence of PAX2 in the samples.

**In vivo tumor grafting**

All procedures involving tumor xenograft mice were conducted in accordance with the guidelines approved by the UT Southwestern Institutional Animal Care and Use Committee (IACUC). Female NOD/scid/gamma mice (Jackson Laboratory) aged 6-8 weeks were used. Ishikawa (ISK) or HEC-1-A cells ($1 \times 10^6$) +/- transduced with *PAX2* lentiviral particles and AN3CA cells ($1 \times 10^6$; both scrambled and *PAX2* knockdown) were subcutaneously injected into the flanks of the mice (n = 4/group) in a total volume of 200 μl consisting of sterile DPBS and Matrigel (1:1). *PAX2* expression was induced by administering doxycycline (Sigma) to mice via drinking water (5 mg/ml supplemented with 5% sucrose) following the injection of ISK and HEC-1-A cells. Tumor dimensions were measured twice weekly using a digital caliper, and tumor volumes were calculated using the formula: (length × width$^2$)/2. Tumor weights were assessed upon completion of the experiment.

**Methylation analysis of *PAX2* from human EC patient samples and cell lines**

*a) <u>Sample collection and DNA preparation</u>*

FFPE tissue samples were endometrial biopsies obtained from 40 patients diagnosed with proliferative endometria. Genomic DNA was extracted from an 8 µm tissue section using the ReliaPrep FFPE gDNA Miniprep System (Promega #A2352) and from human EC cell lines using the QIAamp DNA Mini Kit (Qiagen #51306), following the manufacturer's protocol.

*b) <u>Methylation-specific PCR (MSP) of human</u>* <u>PAX2</u>

DNA was denatured and sulfonated with sodium bisulfite using an EpiTect Bisulfite Kit (Qiagen #59104). In each round of bisulfite treatment, a methylated DNA sample (Zymo Research) was used as a positive control. After bisulfite treatment, unmethylated cytosines were converted into uracil (U), whereas methylated cytosines remained unchanged. Subsequently, the modified DNA was subjected to MSP, using primer pairs designed to specifically amplify either methylated or unmethylated DNA. The methylated and unmethylated primer pair sequences were as follows: methyl specific, 5'-GGGTTTTTTCGTCGAAGTTC-3' (sense) and 5'-ACTAAAACCTCGACTCCCGAT-3' (anti-sense); and unmethyl-specific, 5'-GGTTTTTTTGTTGAAGTTTGG-3' (sense) and 5'-AAAACTAAAACCTCAACTCCCAAT-3' (anti-sense), as described previously (*7*). The thermocycler conditions for PCR were as follows: initial denaturation at 95$^0$C for 10 min, denaturation at 94$^0$C for 30 s, annealing at 59$^0$C for 45 s, and extension at 72$^0$C for 1 min for 39 cycles, followed by a final extension at 72$^0$C for 7 min. The PCR products were subjected to gel electrophoresis on a 2% agarose gel, stained with ethidium bromide, and visualized under UV illumination using a ChemiDoc Imaging System (Bio-Rad Labs).

c*)<u>Targeted methylation sequencing (methyl-seq) and data analysis</u>*

Bisulfite sequencing was conducted on DNA samples extracted from eight human EC cell lines, with four expressing PAX2 (non-silenced) and four lacking PAX2 (silenced). DNA was isolated using the QIAamp DNA Mini Kit (Qiagen) according to the manufacturer's instructions. DNA quality was assessed by agarose gel electrophoresis to detect DNA degradation, and the yield was quantified using a Qubit fluorometer. DNA samples were subjected to Agilent SureSelect$^{XT}$ Methyl-seq (Agilent Technologies) analysis to investigate their DNA methylation status across the entire *PAX2* (GRCh38: chr10: 100,629,868-100,859,867) and *MLH1* loci (GRCh38: chr3: 36,960,880-37,060,879) using a custom-designed capture panel for methyl-seq according to the manufacturer's instructions (design Name: UTSW_MethylSeq_121222, design ID: 3435971, species: *H. sapiens* (UCSC hg38, GRCh38, December 2013). The total probe count was 4120 spanning 379.269 kbp. The DNA samples were fragmented and subjected to end repair and dA-tailing to allow the attachment of methylated adapters. Subsequently, the purified DNA fragments were bisulfite-treated using the EZ DNA Methylation Gold Kit (Zymo Research #D5005). The resulting DNA library was generated by PCR amplification and sequencing was performed using the HiSeq platform.

Paired-end reads were trimmed using Trim Galore (v.0.6.10; https://www.bioinformatics.babraham.ac.uk/projects/trim_galore/) to ensure data quality.

Alignment and methylation calling were conducted using Bismark (v.0.24.0) (*8*), which initially mapped trimmed reads to the human GRCh38 reference genome using Bowtie 2 (2.5.1) (*9*). Subsequently, duplicate reads were removed from the alignment results. Methylation levels on cytosines were determined and CpG island coordinates were extracted from the UCSC table (https://hgdownload.cse.ucsc.edu/goldenpath/hg38/database/cpgIslandExt.txt.gz). Methylation coverage of CpG islands was calculated using BEDTools (v.2.30.0) (*10*). Finally, methylation coverage was visualized using the Integrative Genomics Viewer (IGV, v.2.9.4) (*11, 12*).

**Chromatin Immunoprecipitation assay with sequencing (ChIP-Seq) and data analysis**

Human EC cell lines (Ishikawa: control and *PAX2* re-expressed; AN3CA: scrambled and *PAX2* knockdown) and PDX tumor samples were used for ChIP, followed by sequencing. EC cells were grown to 70-80% confluency, trypsinized, and washed in PBS. PDX tumor samples were finely chopped in PBS containing protease inhibitors and washed. Harvested cells were cross-linked with 1% formaldehyde for 8 min at RT, followed by quenching with 1.25M glycine. The cells were then lysed and sonicated, and the chromatin was immunoprecipitated using the Histone H3K27ac antibody (Active Motif #39133) and the HighCell# ChIP kit with protein G (Diagenode), according to the manufacturer's instructions. The ChIP-processed DNA was reverse-crosslinked and purified using high-salt buffers and phenol-chloroform-isoamyl alcohol (Sigma-Aldrich). DNA was quantified using a Qubit fluorometer (Thermo Fisher Scientific), and the samples were used to construct next-generation sequencing libraries using the KAPA HyperPrep kit (KAPA Biosystems). After appropriate size selection by AMPure XP Beads (Beckman Coulter Life Sciences) and quality checks using a 4200 TapeStation (Agilent), the ChIP-Seq libraries were sequenced using NextSeq 2000 (Illumina).

Raw reads were aligned to the hg19 reference genome using BWA (v0.7.17-r1188) (*13*). Picard (v2.20.3; http://broadinstitute.github.io/picard) was used to remove duplicate reads. Peak detection was carried out with MACS2 (v2.1.0) (*14*) using default parameters with input DNA as a control. Average coverage plots and heatmaps were generated using deepTools (v2.4.2) (*15*). ChIP-Seq peak signals were visualized using Integrated Genomics Viewer (IGV, v2.8.0) (*11*).

**Cleavage Under Targets and Tagmentation (CUT&Tag) Sequencing and data analysis**

CUT&Tag was performed using the CUTANA™ CUT&TAG kit (EpiCypher), following the manufacturer's instructions. Briefly, $5 \times 10^5$ cell nuclei were isolated from each reaction and resuspended in activated concanavalin A-coated beads. The nuclear beads were incubated with primary antibodies (H3K27ac, Active Motif #39133; H3K27me3, EpiCypher #13-0055; H3K4me3, EpiCypher #13-0060; and IgG control, EpiCypher #13-0042) followed by incubation with an anti-rabbit secondary antibody (EpiCypher #13-0047). The antibody-bound nuclear beads were tagged with pAG-Tn5 at RT for 1 h, resuspended in Tagmentation buffer containing 10 mM $MgCl_2$ and incubated at 37°C for 1 h. Tagmentation was stopped by the release of SDS and quenching buffer. The sample was then subjected to PCR using the uniquely barcoded i5 and i7 primers. Sequencing was performed on a NextSeq 2000 (Illumina) sequencing platform in a 50 bp × 2 paired-end mode.

CUT&Tag sequencing reads were mapped to the hg19 genome using Bowtie 2 (v2.1.0) (*16*). Duplicate reads were removed using Picard (v2.20.3) and reads with low mapping quality (MAPQ < 30) were excluded using Samtools (v1.9) (*17*). Peak calling was conducted using SEACR (1.3)

(*18*) with IgG as the control. DeepTools (v2.4.2) was used to generate average coverage plots. CUT&Tag peak signals were visualized using the Integrated Genomics Viewer (IGV, v2.8.0) (*11*).

**Assay for Transposase-Accessible Chromatin with Sequencing (ATAC-Seq)**

To analyze the chromatin encompassing the *PAX2* region, ATAC-seq was performed on PDX tumor samples using established protocols (*19*). Tumors were finely chopped, washed, and lysed using ice-cold lysis buffer (10 mM Tris-HCl pH 7.5, 10 mM NaCl, 3 mM $MgCl_2$, 0.1% NP40, 0.1% Tween-20, and 0.01% digitonin). Nuclei were washed again in cold lysis buffer without NP40 and Digitonin and then incubated with Tn5 transposase mixture (2.5μl of Tn5, 1% digitonin, 10% Tween-20 in 1x TD buffer) at $37^0$C for 30 min in a Thermomixer (Eppendorf). DNA was purified using the DNA Clean and Concentrator-5 kit (Zymo Research) and used for PCR with Nextera adapters. The final product was purified and sequenced using NextSeq 2000 (Illumina).

ATAC-Seq reads were aligned to the hg19 genome using the Bowtie 2 software (v2.1.0). To ensure data quality, duplicate reads were removed using Picard (v2.20.3) and reads with low mapping quality (MAPQ < 10) were excluded using Samtools (v1.9). Peaks were identified using MACS2 (v2.1.0). Average coverage plots were generated using DeepTools (v2.4.2).

**RAD21 Chromatin Interaction Analysis with Paired-End Tag (ChIA-PET) Sequencing**

RAD21 ChIA-PET data were obtained from the ENCODE project (*20*). Looping interactions were identified using ChIA-PET2 (v0.9.3) (*21*).

**RAD21 Chromatin Interaction Predictor (ChIPr) analysis**

Publicly available RAD21 ChIP-seq peaks for the Ishikawa cell line (GEO accession: GSM1010801) were used for ChIPr analysis. The analysis focused on the peaks surrounding the *PAX2* gene and its neighborhood. The strength of all pairwise RAD21-mediated interactions between these peaks was predicted using ChIPr (*22*). To identify the strongest interactions, the loops with interaction strength cutoffs ranging from '3' to '10' were visualized.

**Total RNA isolation, cDNA synthesis and quantitative RT-PCR**

For quantitative PCR analysis, total RNA was isolated from control or *PAX2* re-expressed EC cell lines and from wild-type or *Pax2* cKO organoids using the RNeasy Plus Mini kit (Qiagen #74136), following the manufacturer's protocol. RNA quality and concentration were determined using a NanoDrop 2000 spectrophotometer (Thermo Fisher Scientific). Subsequently, 1 μg of RNA was reverse-transcribed into cDNA using SuperScript VILO Master Mix (Thermo Fisher Scientific). The resulting cDNA was then subjected to amplification using the following sequence-specific primers: human *PAX2*:5'-CCCAGCGTCTCTTCCATCA-3' (sense), 5'-GGCGTTGGGTGGAAAGG-3' (anti-sense); human *GAPDH*: 5'-GCCACATCGCTCAGACACCAT-3' (sense), 5'-GAAGGGGTCATTGATGGCAA-3' (antisense); mouse *Pax2*:5'-AAGCGACAGAACCCGACTATGT-3' (sense), 5'-ACTCCTGTCCCTGCCCCAT-3' (anti-sense); mouse *Gapdh*: 5'-CAACTACATGGTCTACATGTT-3' (sense), 5'-CTCGCTCCTGGAAGATG-3' (anti-sense). Quantitative real-time PCR (qRT-PCR) was performed to analyze the expression levels of *PAX2* and *GAPDH* genes using the $RT^2$ SYBR Green ROX qPCR Mastermix (Qiagen) on a QuantStudio Real-Time PCR System (Applied Biosystems). The qRT-PCR protocol involved a pre-incubation step, followed by 40 amplification cycles. The comparative delta-delta Ct ($2^{-\Delta\Delta Ct}$) method was used to evaluate the expression level of the target gene relative to the housekeeping gene (*GAPDH*) across different experimental groups.

**Library preparation, RNA-Seq and data analysis**

Total RNA samples from Ishikawa cells (both control and *PAX2* re-expressed) were submitted to the McDermott Next Generation Sequencing Core at UT Southwestern for library preparation. 1 µg of DNase-treated high-quality RNA (RIN score ≥ 8 as determined by the Agilent 2100 Bioanalyzer) was prepared using the TruSeq Stranded Total RNA LT Sample Prep Kit (Illumina). Total RNA was depleted of rRNA and fragmented prior to strand-specific cDNA synthesis. The cDNA was poly A-tailed and ligated with indexed adapters. Following adapter ligation, the samples were PCR-amplified, purified with Ampure XP beads, and revalidated using the Agilent 2100 Bioanalyzer. The samples were quantified using a Qubit before normalization and run on an Illumina HiSeq 2500 using SBS v3 reagents. The samples were sequenced on the Illumina NextSeq 500 with a read configuration of 75 bp and single end reads, generating 25-35 million reads per sample.

The quality of raw RNA sequencing data was assessed using FastQC (v0.11.8) (*23*). The reads were aligned to the hg19 reference genome using STAR aligner (v2.7.3a) (*24*). The gene expression levels for each sample were quantified using RSEM (v1.3.2) (*25*)with default parameters. Differentially expressed genes were identified using DESeq2 (v1.36.0) (*26*) by applying a threshold of $\log_2 FC > 1$ and $FDR < 0.01$.

**Gene ontology analysis**

Gene ontology analyses were performed through the Gene Ontology Resource using the PANTHER Overrepresentation Test with all *Homo sapiens* genes in the database (*27, 28*). P-values were determined using Fisher's Exact Test with False Discovery Rate Correction.

**Single-cell RNA Sequencing (scRNA-Seq) and data analysis**

*a) Sample preparation*

Uterine horns from *Pax2* cKO mice were dissected immediately after euthanasia, washed in $Ca2^+$- and $Mg2^+$-free Dulbecco's phosphate-buffered saline (DPBS), and minced into 2-3 mm longitudinal pieces using a sterile scalpel. The minced tissue fragments were digested in a cocktail of Collagenase V (0.4 mg/ml, Sigma) and Dispase II (1.25 U/ml, Sigma) at $37^0$C for 1 h with intermittent shaking. Digestion was stopped by adding Advanced DMEM/F12 medium supplemented with 10% FBS, and cell debris was removed using a 100 µm cell strainer (Fisher Scientific). The isolated uterine cell suspension was centrifuged at 700 × g for 5 min and the cell pellet was resuspended in PBS containing 0.4% BSA. Cell viability was checked using a Countess II Automated Cell Counter (Invitrogen). Cells were then processed in the Chromium Single Cell Gene Expression platform (10x Genomics) to build the 3′ transcriptome libraries and sequenced at the McDermott Next Generation Sequencing Core at UT Southwestern Medical Center.

*b) Analyses of single-cell transcriptomes*

The mkfastq module of Cell Ranger (v5.0.1, 10X Genomics) was used to convert BCL files into the FASTQ format. Next, the Cell Ranger count module was used to align reads from each library to the mouse reference genome (mm10), and transcript counts of each cell were quantified using unique molecular identifiers (UMIs) and valid cell barcodes. Cell Ranger used the EmptyDrops

(*29*) method to identify populations of cells with low RNA content. Briefly, this algorithm first applies a threshold based on the total UMI counts per barcode to identify cells, distinguishing the initial set of cells with high RNA content. We then examined the RNA profiles of the remaining barcodes to differentiate between empty and cell-containing partitions. Cell Ranger's graph-based clustering method was used to cluster the cells into different cell types. Differentially expressed genes between the defined clusters were identified using Cell Ranger differential expression analysis.

**Endometrial organoids**

*a) Uterine epithelial cell isolation, organoid culture, and expansion*

Endometrial organoids were generated from epithelial cells isolated from 12-16 weeks old control, WT, *Pax2*, *Pten*, and *Pax2*/*Pten* cKO mice, as described previously (*30*) with slight modifications. The dissected uteri were washed in $Ca2^{+}$-and $Mg2^{+}$-free Dulbecco's phosphate-buffered saline (DPBS; Gibco) and sliced into 2-3 mm longitudinal pieces using a sterile scalpel. Subsequently, the tissue fragments were digested in a solution containing Collagenase V/Dispase II/Trypsin enzymes (0.4 mg/ml Collagenase V, 1.25 U/ml Dispase II, and 0.05% trypsin-EDTA; Sigma) diluted in Advanced DMEM/F12 medium (Thermo Fisher Scientific) at $37^{0}$C for 1 h with intermittent shaking. The digestion process was stopped by adding Advanced DMEM/F12 medium supplemented with 10% FBS, and the digestion mixture was triturated several times to disrupt the epithelial glands. The resulting mixture was filtered through a 100 µm cell strainer to remove any cell debris, and the filtrate was collected and passed through a 20 µm cell strainer to separate the stromal cells. The epithelial cells were isolated by turning the 20 µm cell strainer upside down and washing the epithelial cells into a sterile 50 ml tube using 10 ml of HBSS (Gibco) supplemented with 10% FBS.

Isolated epithelial cells were pelleted by centrifugation at 700 × g for 5 min and resuspended in Matrigel (Corning). Subsequently, 20 µL of Matrigel-cell suspension (approximately 500 cells) was seeded into individual wells of a 48-well plate and allowed to solidify at 37°C for 5 min. The cell-containing dome was overlaid with 250 µL of organoid expansion medium, which consisted of Advanced DMEM/F12 medium (Gibco) supplemented with 1% penicillin/streptomycin, 2mM Glutamax (Thermo Fisher Scientific), 1 mM nicotinamide (Sigma), 1% insulin-transferrin-selenium (Thermo Fisher Scientific), 1% N-2 supplement (Thermo Fisher Scientific), 2% B-27 supplement (Thermo Fisher Scientific), 100 ng/ml R-Spondin1 (Peprotech), 100 ng/ml Noggin (Peprotech), 100 ng/ml FGF10 (R&D Systems), 50 ng/ml EGF (Peprotech), and 50 ng/ml Wnt3a (Peprotech).

The organoids were maintained in culture by refreshing the organoid expansion medium every 3 days, and passaging was conducted every 5-7 days. For passaging, organoids were dissociated into single cells by incubating them in TrypLE Express (Thermo Fisher Scientific) or 0.05% Trypsin-EDTA (Gibco) at $37^{0}$C for 10 min. Trypsinization was stopped by adding DMEM/F-12 supplemented with 10% FBS. Subsequently, cell clumps were further dissociated into single cells by gentle pipetting and reseeded in Matrigel droplets (500 cells/20µl droplet).

*b) Harvesting, fixing, and staining of the organoids*

After 5-7 days of culture, the organoids were harvested by dissolving the Matrigel dome in ice-cold cell recovery solution (Corning) for 30 min at $4^{0}$C. Following this, the harvested organoids were washed in PBS, and fixed in 4% PFA (Electron Microscopy Sciences) for 20 min at RT. Once fixed, the organoids were washed again in PBS, gently pelleted, overlaid with 200 µL of low-melting agarose (Lonza) at $42^0$C, and allowed to solidify. Subsequently, the agarose plugs containing organoids were embedded in paraffin, sectioned at 5 µm thickness, and stained with H&E. For immunostaining, the sections were deparaffinized in xylene and rehydrated using a graded ethanol series. Antigen retrieval was performed using a sodium citrate buffer (pH 6.0, Vector Laboratories). The slides were washed with TBS containing 0.1% Tween 20 (Sigma) and incubated overnight at 4°C with primary antibodies against PAX2 (Sigma #311R-1, 1:100) and pan-cytokeratin (Abcam #ab7753, 1:200). Organoids were stained with phalloidin (Sigma #P5282, 1:400) to stain actin filaments. After washing, the sections were incubated with Alexa Fluor secondary antibodies (Alexa Fluor 555 goat anti-rabbit IgG, Thermo Fisher Scientific #A-21429, 1:250 or Alexa Fluor 488 goat anti-mouse IgG, Thermo Fisher Scientific #A-11029, 1:250) for 1 h at RT and coverslips were mounted using VECTASHIELD antifade mounting medium with DAPI (Vector Laboratories). Fluorescent images were taken on a Nikon ECLIPSE Ni fluorescence microscope. For 3D z-stack imaging, whole-mount organoids were fixed, permeabilized, stained with antibodies, and imaged using a Zeiss LSM880 inverted confocal microscope.

*c) Organoid proliferation assay*

To assess organoid growth rates, epithelial cells from control, *Pax2*, *Pten*, and *Pax2*/*Pten* cKO mice were mixed with Matrigel and 500 cells were seeded in triplicate into the wells of a 96-well plate using 20 µL Matrigel drops. The Matrigel-domed cells were then overlaid with 100 µL of organoid expansion medium and incubated at 37°C with 5% $CO_2$. ATP luminescence, indicative of cell viability, was measured on days 0, 2, 4, 6, and 8 by adding 100 µL of Cell Titer-Glo 3D reagent (Promega #G9683) to actively growing organoids and recording luminescence using a Tecan Spark 10M microplate reader. Every other day, organoids were imaged using a phase-contrast microscope (Olympus). The diameter of the organoids was quantified using ImageJ software (NIH), allowing for a comparison of sizes among different organoids.

**Histology, immunohistochemistry (IHC), and quantification**

Normal uteri or uterine tumor tissues from control, *Pax2*, *Pten*, and *Pax2*/*Pten* cKO mice were fixed overnight in 10% buffered formalin at 4°C. Subsequently, the tissues were processed for paraffin embedding, sectioned at 5 µm thickness, and stained with H&E. H&E images were captured using a DS-Ri2 phase-contrast microscope (Nikon, Tokyo, Japan). For immunostaining, mouse tissue sections and human EC TMA slides (US Biomax) were subjected to deparaffinization in xylene, followed by rehydration in graded alcohol and water. Antigen retrieval was performed using sodium citrate buffer (pH 6.0, Vector Laboratories). The slides were washed with TBS containing 0.1% Tween 20 (Sigma) and incubated with 3% hydrogen peroxide ($H_2O_2$; Fisher Scientific) in water for 15 min to block endogenous peroxidase activity. The tissue sections were subsequently blocked with BLOXALL endogenous peroxidase and alkaline phosphatase blocking solution (Vector Laboratories) for 10 min at RT. Following blocking, tissue sections were incubated with primary antibodies against PAX2 (Sigma #311R-1, 1:50), PTEN (Cell Signaling Technology #9559, 1:200), or p63 (GeneTex #GTX102425, 1:1000) overnight at 4°C. After incubation, the slides were washed and incubated with peroxidase-conjugated secondary antibodies (Vector Laboratories #MP-7401) followed by DAB substrate (Agilent Dako #K346811-2) to detect the bound antibodies. Tissue sections were counterstained with hematoxylin and mounted using Permount mounting medium (Fisher Scientific). Images were acquired using an

Aperio AT2 slide scanner (Leica Biosystems) and subsequently analyzed using the Aperio ImageScope software. The IHC intensity score (H-score) was calculated from pixel intensity values, which was the sum of three times the percentage of pixels with strong staining, 2 times the percentage of pixels with moderate staining, and 1 time the percentage of pixels with weak staining.

**RNA in situ hybridization (ISH)**

*PAX2* RNA ISH was performed on human endometrial FFPE tissue sections using the RNAscope 2.5 HD Assay-BROWN kit (Advanced Cell Diagnostics #322300) and the RNAscope probe Hs-*PAX2* (Advanced Cell Diagnostics #442541), following the manufacturer's protocols. Images were captured using a Nikon DS-Ri2 phase-contrast microscope.

**Fluorescence in situ hybridization (FISH)**

FISH was conducted on 4 µm tissue sections from 12 randomly selected cases of endometrioid intraepithelial neoplasia (EIN) identified as PAX2-silenced by IHC. Dual-color break-apart *PAX2* probes (Empire Genomics #PAX2BA-20-ORGR) specific for the 5′ and 3′ regions of 10q24.31 were utilized. The 5′ fragment (~162 kb) of the probe was labeled with TAMRA (orange signal), whereas the 3′ fragment (~188 kb) was labeled with fluorescein (green signal). Probe labeling and hybridization were performed according to the manufacturer’s instructions (*31*). FISH images were acquired on a Zeiss axioimager M2 scope (100X objective) with applied spectral imaging software.

**Immunofluorescence (IF)**

EC cell lines were cultured on 8-well chambered glass slides and fixed in 4% paraformaldehyde (Electron Microscopy Sciences) for 20 min before processing for IF. The fixed cells were washed in PBS and permeabilized in PBS containing 0.5% Triton X-100 (Sigma) for 10 min at 4°C, followed by rinsing in PBS/glycine buffer (PBS containing 0.1 M Glycine). Subsequently, the cells were blocked in IF buffer (130 mM NaCl, 7 mM $Na_2HPO_4$, 3.5 mM $NaH_2PO_4$, 0.1% BSA, 0.2% Triton X-100, and 0.05% Tween 20) supplemented with 10% goat serum (Vector Laboratories) for 1 h and then incubated with primary antibody against PAX2 (Sigma #311R-1, 1:100) overnight at 4°C. To stain mouse uterine tissue sections, the slides were first deparaffinized, followed by antigen retrieval using sodium citrate buffer (pH 6.0, Vector Laboratories). Tissue sections were incubated in blocking buffer (TBS containing 0.1% Triton X-100 and 5% goat serum) for 1 h at RT and then incubated with primary antibodies against ERα (Cell Signaling Technology #13258, 1:200) and PR-A/B (Cell Signaling Technology #8757, 1:200) overnight at 4°C. The next day, the slides were washed three times in TBS (Vector Laboratories) containing 0.1% Tween 20 for 5 min each and incubated with Alexa Fluor secondary antibodies for 1 h at RT. Finally, the slides were cover-slipped using VECTASHIELD antifade mounting media with DAPI (Vector Laboratories), and images were acquired using a Nikon ECLIPSE Ni fluorescence microscope with NIS-Elements BR software or a Zeiss LSM880 inverted confocal microscope.

**Western blotting**

Cultured EC cells were washed with PBS and lysed in ice-cold RIPA buffer (Thermo Fisher Scientific) for 1 h on ice supplemented with protease and phosphatase inhibitors (Sigma). The cell lysates were centrifuged for 5 min at 13,000 rpm and 4°C, and the resulting supernatants were collected. The protein concentration of each sample was determined using a Pierce BCA Protein

Assay Kit (Thermo Fisher Scientific #23225). 25 µg of protein was mixed with 4X NuPAGE LDS Sample Buffer (Thermo Fisher Scientific), heated at $95^0$C for 5 min, and resolved on 4-12% NuPAGE gels (Thermo Fisher Scientific) at 120 V for 1.5 hrs. Proteins were transferred to a polyvinylidene fluoride (PVDF) membrane (Sigma) using the Trans-Blot Turbo Transfer System (Bio-Rad Labs). After transfer, the membrane was blocked in TBS-T (0.1% Tween 20 in TBS) containing 5% skim milk (w/v) for 1 h at RT and then probed with primary antibodies against PAX2 (Sigma #311R-1, 1:1000), MLH1 (Abcam #ab92312, 1:1000), HIF1AN (Abcam #ab92498, 1:1000), GAPDH (Cell Signaling Technology #2118, 1:4000), or ACTIN (Cell Signaling Technology #3700, 1:4000) overnight at 4°C. The membrane was washed and incubated with a horseradish peroxidase (HRP)-conjugated secondary antibody (donkey anti-rabbit IgG, HRP linked Ab, Amersham #NA934, 1:2000 or sheep anti-mouse IgG, HRP-linked Ab, Amersham #NA931, 1:2000) for 1 h at RT and developed using SuperSignal West Dura Extended Duration Substrate (Thermo Fisher Scientific #34076) for HRP detection. Protein bands were visualized by chemiluminescence using a ChemiDoc Imaging System (Bio-Rad Labs). β-Actin or GAPDH served as loading control.

**Real-time cell proliferation and scratch wound assay by IncuCyte live-cell imaging**

*PAX2* +/- Ishikawa, HEC-1-A, and HEC-1-B cells were seeded at a density of 5,000 cells/well in a flat-bottom 96-well plate and incubated at 37°C in a humidified $CO_2$ incubator. Six hours later, the plate was transferred to an IncuCyte live-cell imaging system (Essen Bioscience). The cells were allowed to grow until reaching 90-100% confluency, and bright-field images were captured at five fields per well every 6 h. Cell proliferation curves were generated using IncuCyte software by plotting normalized cell confluency, which represents the ratio of confluency at 6 h intervals to confluency at the start of the experiment.

For wound closure assays, 5,000 AN3CA cells (both scrambled and *PAX2* knockdown) were seeded in a 96-well microplate and cultured to reach 90-100% confluency. Subsequently, a single homogenous and reproducible scratch was created per well using the WoundMaker tool (Essen Biosciences). Following scratching, the cells were washed twice with PBS to remove any detached cells from the scratched area. After washing, 150 µL of MEM containing 10% FBS was added to each well, and the plate was then placed into the IncuCyte live-cell imaging system. Real-time images of wound closure were captured every 3 h, and the wound width (distance between leading edges of each cell front) was calculated using the IncuCyte software. The percentage of wound closure was determined by analyzing the ratio of wound width at 3 h intervals to wound width at time 0.

**Cell cycle analysis**

AN3CA cells, including both scrambled and *PAX2* knockdown variants, were synchronized in the G1 phase by inducing growth arrest via serum starvation for 6 h. Subsequently, the starved cells were cultured in complete medium until they reached a confluency–70-80%. The subconfluent cells were then fixed in 70% alcohol solution for 10 min at RT. Fixed cells were washed with PBS and permeabilized in PBS containing 0.1% Triton X-100 for 15 min at RT. After permeabilization, the cells were passed through a 40 µm nylon cell strainer (Fisher Scientific) to remove any cell debris. Following this, the cells were again washed with PBS and incubated in 50 µg/mL propidium iodide (PI, BD Biosciences #556463) to stain DNA and 100 µg/mL RNase A (Thermo Fisher Scientific) to remove excess RNA. Approximately $1 \times 10^6$ cells were analyzed using a FACS analyzer equipped with a 488 nm LASER line to detect PI-stained nuclei. The percentage of cells in the G0/G1, S, and G2/M phases of the cell cycle was determined by analyzing the histogram output using FlowJo software.

**Clonogenic assay**

The disparity in colony-forming capability between Ishikawa cells (both ISK-pLVX and ISK-pLVX-*PAX2*) was evaluated by seeding 2,000 cells/well in a 6-well plate and incubating them in complete medium for 5-7 days until colonies formed. On day 7, cells were fixed in 70% alcohol and stained with 0.2% crystal violet (Sigma-Aldrich). Colonies, defined as groups of 50 or more cells, were quantified using ImageJ software.

**Statistics**

For all experiments, data are presented as the mean ± SEM unless otherwise stated, and statistical analyses were performed using GraphPad Prism (v.10.0.0). Fisher's exact test was used to determine the P-value for PAX2 loss between young and aged women. PAX2 H-scores among different grades of patients were determined using the Mann-Whitney U test. Statistical significance between two groups or multiple groups was assessed by unpaired, 2 tailed Student's *t* test and one-way ANOVA, respectively. Kaplan-Meier analysis and the log-rank test were used to determine the P value between survival curves. Each in vitro experiment was repeated thrice, and the in vivo experiments were repeated twice. Statistical significance was set at $P < 0.05$.

### Supplementary Figures 1 to 7

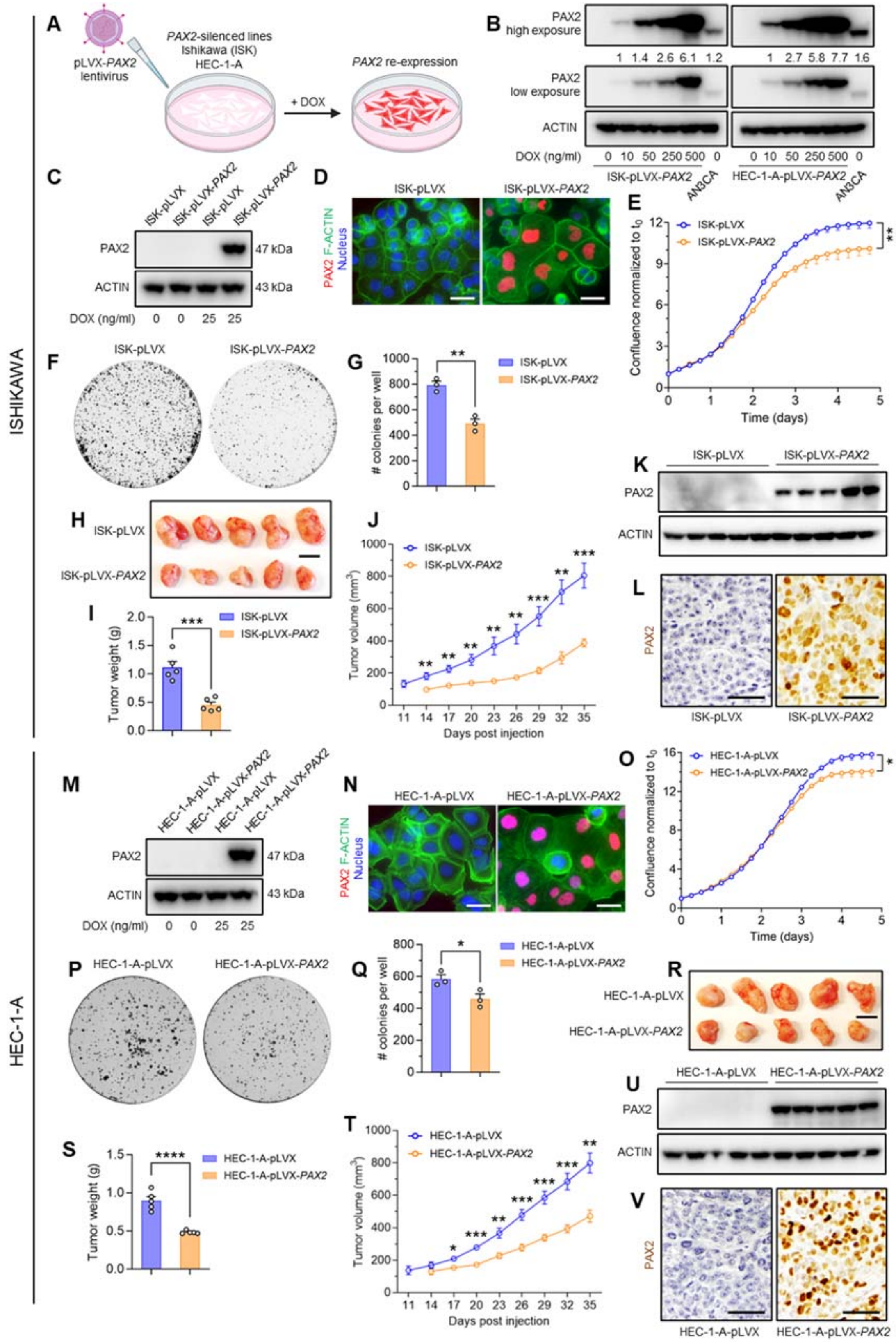

**Supplementary Fig. 1.** ***PAX2*** **re-expression in PAX2-silenced lines suppresses cell growth.**

(**A**) Schematic of experimental approach (created with BioRender.com).
(**B**) Western blot analysis of PAX2 expression following lentiviral transduction in Ishikawa (ISK) ISK-pLVX-*PAX2* and HEC-1-A-pLVX-*PAX2* cells after doxycycline (DOX) induction. PAX2 levels were correlated with [DOX]. AN3CA was used as the control. The lower apparent PAX2 molecular weight in AN3CA is attributed to the small size of PAX2 and the C-terminal Myc-DDK-tag.
(**C**) Western blot analysis of Ishikawa cells (ISK) following lentiviral transduction with an empty vector (ISK-pLVX) or pLVX-*PAX2* (ISK-pLVX-*PAX2*). Cells were cultured with +/- 25 ng/mL DOX for 24h.
(**D**) Immunofluorescence confirmed nuclear localization of re-expressed PAX2 (red). F-actin and DAPI counterstaining highlighted cell membranes (green) and nuclei (blue). Bars=50 µm.
(**E**) Live cell image analysis showing *PAX2*-mediated cell growth suppression; proliferation is expressed as normalized confluency to $t_0$ (n=3, mean±SEM, unpaired 2-tailed t test).
(**F**) Clonogenic assay, representative images of ISK-pLVX and ISK-pLVX-*PAX2* cells stained with crystal violet following seeding of equal numbers of cells.
(**G**) Clonogenic assay and quantitative colony counts (n=3, mean±SEM, unpaired 2-tailed t test).
**(H)** Xenografts of ISK-pLVX and ISK-pLVX-*PAX2* cells in flanks of NOD/scid/gamma females (n=5) following injection of $10^6$ cells. Tumors were harvested 35 days after the subcutaneous injection. Bar=1 cm.
**(I)** Tumor weights on day 35 (n=5, mean±SEM, unpaired 2-tailed t test).
(**J**) Tumor volume of ISK-pLVX and ISK-pLVX-*PAX2* xenografts per caliper measurement (n=5, mean±SEM, unpaired 2-tailed t test).
(**K**) Western blot analysis of lysates from ISK-pLVX and ISK-pLVX-*PAX2* xenografts at harvest, confirming their stable expression.
(**L**) PAX2 immunostaining of ISK-pLVX and ISK-pLVX-*PAX2* xenografts, confirming stable expression and nuclear localization. Bars=50 µm.
(**M**) Western blot analysis validated that 25 ng/mL DOX led to enforced PAX2 expression in HEC-1-A cells following lentiviral transduction (HEC-1-A-pLVX-*PAX2*). Lentiviral transduction with empty vector (HEC-1-A-pLVX) was used as the negative control.
(**N**) Immunofluorescence staining reveals anticipated nuclear localization of PAX2 (red) in HEC-1-A-pLVX-*PAX2* cells, DAPI counterstains nuclei (blue). Bars=50 µm.
(**O**) Cell proliferation assays using live cell imaging of HEC-1-A-pLVX and HEC-1-A-pLVX-*PAX2* cells showed growth suppression by PAX2. % confluency was normalized to $t_0$ (n=3, mean±SEM, unpaired 2-tailed t test).
(**P**) Clonogenic assay: representative images of HEC-1-A-pLVX and HEC-1-A-pLVX-*PAX2* cells stained with crystal violet.
(**Q**) Colony number (n=3, mean±SEM, unpaired 2-tailed t test).

(**R**) Xenografts of HEC-1-A-pLVX and HEC-1-A-pLVX-*PAX2* cells in the flanks of NOD/scid/gamma female mice (n=5). Tumors were harvested 35 days after the subcutaneous injection. Bar=1 cm.
(**S**) Tumor weights on day 35 (n=5, mean±SEM, unpaired 2-tailed Student's t-test).
(**T**) Tumor volume growth of HEC-1-A-pLVX and HEC-1-A-pLVX-*PAX2* xenografts per caliper measurement (n=5, mean±SEM, unpaired 2-tailed t test).
(**U**) Western blot analysis of PAX2 protein expression in lysates from HEC-1-A-pLVX and HEC-1-A-pLVX-*PAX2* xenograft tumors.
(**V**) PAX2 immunolocalization in tumor sections from HEC-1-A-pLVX and HEC-1-A-pLVX-*PAX2* xenografts. Bars=50 μm.
For all panels, *P<0.05; **P<0.01; ***P<0.001; ****P<0.0001.

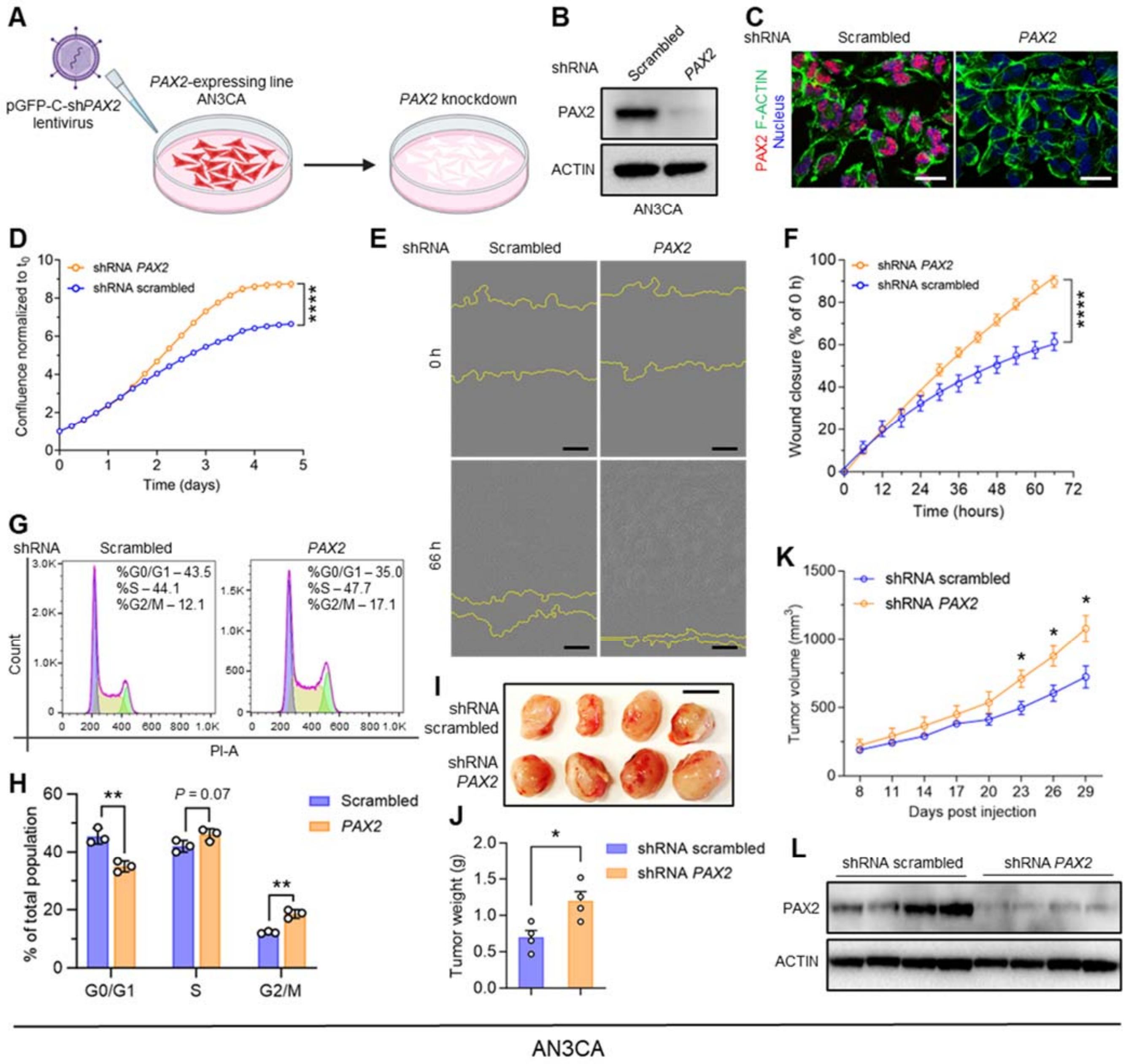


**Supplementary Fig. 2. *PAX2* knockdown (KD) in PAX2-expressing line AN3CA promotes cell growth.**

(**A**) Schematic of experimental approach (created with BioRender.com).

(**B**) Western blot confirmation of lentiviral shRNA-mediated *PAX2* KD in AN3CA cells.

(**C**) Immunofluorescence of PAX2 (red) demonstrating expected reduction in nuclear PAX2 in *PAX2* KD AN3CA cells. F-actin and DAPI counterstaining highlighted cell membranes (green) and nuclei (blue). Bars=50 µm.

(**D**) Live cell analysis shows *PAX2* KD-mediated cell proliferation in AN3CA cells compared to scrambled shRNA control; confluency normalized to $t_0$ (n=3, mean±SEM, unpaired 2-tailed t test).

(**E**) Wound healing assay and representative images of AN3CA cells (scrambled vs. *PAX2* KD) at $t_0$ and 60h. Bars=250 μm.
(**F**) Wound healing assay and continuous live analysis (n=3, mean±SEM; unpaired 2-tailed t test).
(**G**) Cell cycle analysis of AN3CA cells (scrambled vs. *PAX2* KD, n=3) by propidium iodide flow cytometry. Peaks in purple, yellow, and green indicate the percentage of cells in G0/G1, S, and G2/M phases, respectively.
(**H**) Quantitative analysis of G0/G1, S, and G2/M phases (n=3, mean±SEM, multiple 2-tailed t tests).
(**I**) Xenografts of AN3CA cells (scrambled vs. *PAX2* KD) in flanks of NOD/scid/gamma female mice (n=4). Bar=1 cm.
(**J**) Tumor weight at harvest (day 29, n=4, mean±SEM, unpaired 2-tailed t test).
(**K**) Tumor volume growth of AN3CA (scrambled vs. *PAX2* KD) xenografts per caliper measurement (n=4, mean±SEM, unpaired 2-tailed t test).
(**L**) Western blotting of lysates from AN3CA (scrambled vs. *PAX2* KD) xenografts at end of experiment, confirming stable KD.
For all panels, *P<0.05 **P<0.01; ****P<0.0001.

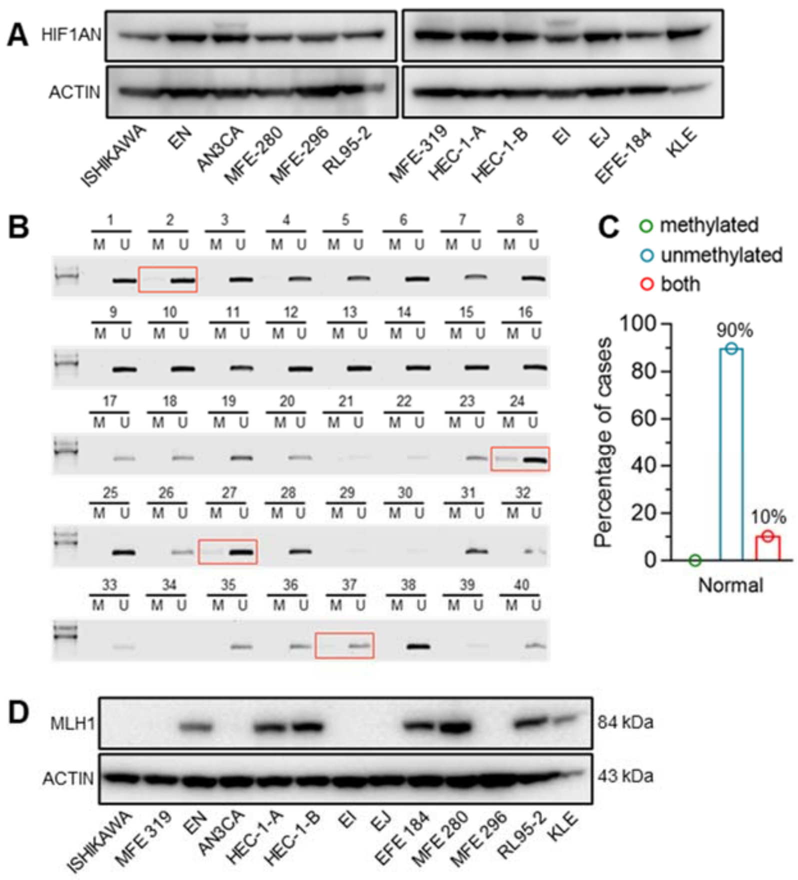


**Supplementary Fig. 3. Ancillary data for Fig. 2 including demonstration that *PAX2* locus is unmethylated in normal endometria.**

(**A**) Western blot analysis of protein of neighboring gene (*HIF1AN*) confirming lack of silencing in an expanded set of EC lines.

(**B**) Methylation-specific PCR for *PAX2* locus using methyl-specific (M) and unmethyl-specific (U) oligonucleotide primers was performed on bisulfite-treated normal human endometrial samples (n=40). The methylation-specific PCR assay used the same primers and conditions as in (*7*). Rare samples with partial methylation are indicated by rectangles. The results show that *PAX2* is unmethylated in normal endometria, contrary to a prior report (*7*).

(**C**) Fraction of cases yielding MS-PCR products corresponding to the methylated/unmethylated state or both.

(**D**) Western blot analysis confirmed the loss of protein in three EC lines with *MLH1* CpG island hypermethylation. Ishikawa is known to have a high mutational burden (2745 mutations) and to be MSI-high (*32*). Using the NGS platform for EC genes (*33*), we identified biallelic pathogenic *MLH1* mutations in known hotspots *c.1731+5G>T* and *c.790+3delA*, showing that these mutations inactivate MLH1 (*34*). MFE-319 and MFE-296 likely exhibit *MLH1* promoter hypermethylation (not tested).

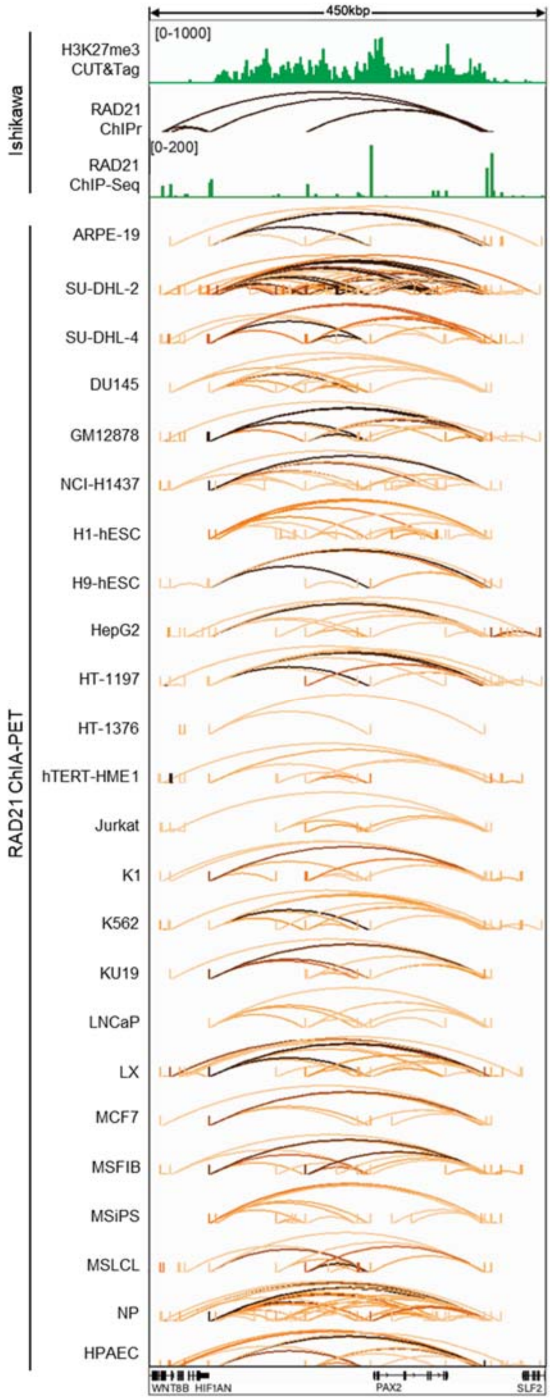


**Supplementary Fig. 4. RAD21 ChIA-PET across 24 cell lines.**

The analysis focused on the H3K27me3 enriched region surrounding *PAX2*, which was identified as a highly conserved insulated neighborhood. This neighborhood is bound by RAD21-mediated chromatin loops, highlighting RAD21's role in regulating chromatin architecture within this region.

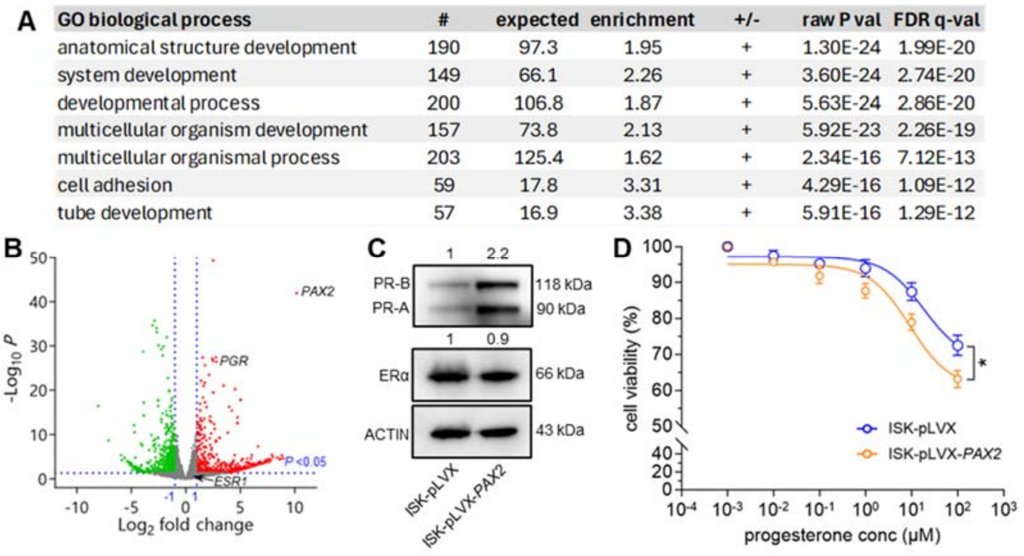

| GO biological process | # | expected | enrichment | +/- | raw P val | FDR q-val |
|---|---|---|---|---|---|---|
| anatomical structure development | 190 | 97.3 | 1.95 | + | 1.30E-24 | 1.99E-20 |
| system development | 149 | 66.1 | 2.26 | + | 3.60E-24 | 2.74E-20 |
| developmental process | 200 | 106.8 | 1.87 | + | 5.63E-24 | 2.86E-20 |
| multicellular organism development | 157 | 73.8 | 2.13 | + | 5.92E-23 | 2.26E-19 |
| multicellular organismal process | 203 | 125.4 | 1.62 | + | 2.34E-16 | 7.12E-13 |
| cell adhesion | 59 | 17.8 | 3.31 | + | 4.29E-16 | 1.09E-12 |
| tube development | 57 | 16.9 | 3.38 | + | 5.91E-16 | 1.29E-12 |



**Supplementary Fig. 5.** ***PAX2*** **regulates** ***PGR*** **among large number of genes involved in developmental processes.**

(**A**) Gene ontology analysis of DEGs.

(**B**) Volcano plot showing DEGs (P<0.05) in ISK-pLVX-*PAX2* cells compared with the empty vector control (ISK-pLVX). Vertical lines indicate $\log_2$ fold change threshold of ±1 and dotted horizontal line marks a P-value threshold of 0.05.

(**C**) Western blot analysis of estrogen receptor α (ERα) and progesterone receptor A/B (PR-A/B) in ISK-pLVX and ISK-pLVX-*PAX2* cells.

(**D**) Cell survival assay showing modest growth reduction following progesterone treatment in *PAX2*-expressing Ishikawa cells (n=3). Data are shown as the mean±SEM; 2-tailed t test, *P<0.05.

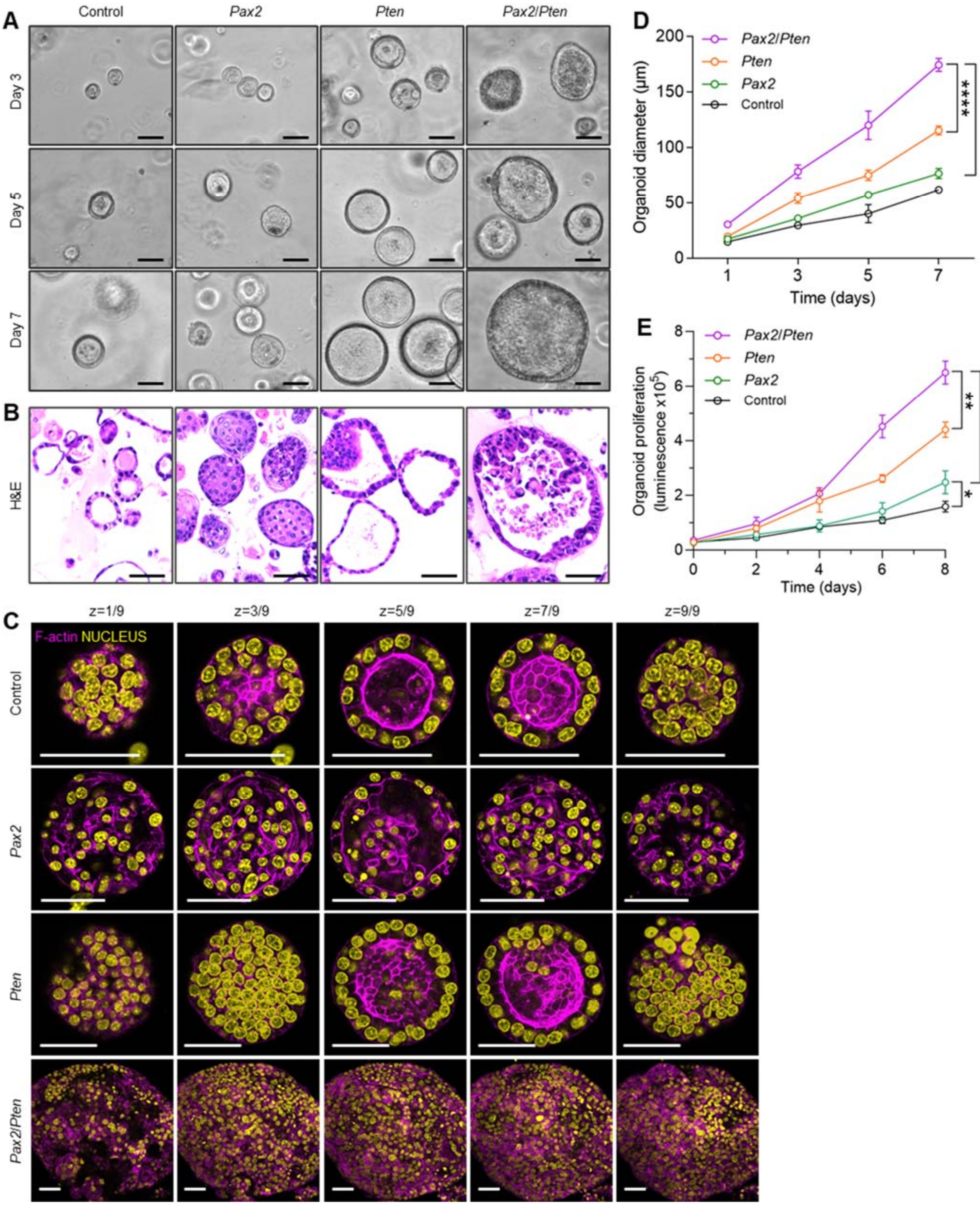
A
Control
Pax2
Pten
Pax2/Pten
Day 3
Day 5
Day 7
B
H&E
C
z=1/9
z=3/9
z=5/9
z=7/9
z=9/9
F-actin NUCLEUS
Control
Pax2
Pten
Pax2/Pten
D
Organoid diameter (μm)
Time (days)
Pax2/Pten
Pten
Pax2
Control
E
Organoid proliferation (luminescence x10^5)
Time (days)
Pax2/Pten
Pten
Pax2
Control

**Supplementary Fig. 6. Studies of organoids from *Pax2*, *Pten*, and *Pax2/Pten* endometria show synergistic growth and morphologic phenotypes.** Organoids harvested from mice at 12-16 weeks of age (before onset of neoplasia).

(**A**) Phase-contrast images of endometrial epithelial organoids at days 3, 5, and 7 after harvesting from uteri. Bars=50 μm.

(**B**) H&E-stained sections from paraffin-embedded formalin-fixed organoids. Bars=50 μm.

(**C**) Confocal 2 μm slices of organoids stained with F-actin (magenta) and DAPI (yellow). Bars=50 μm.

(**D-E**) Comparison of organoid diameter (D) and cell number (E) by ATP luminescence among the four genotypes (n=3, mean±SEM). *P<0.05, **P<0.01; ***P<0.001; ****P<0.0001, unpaired 2-tailed t test.

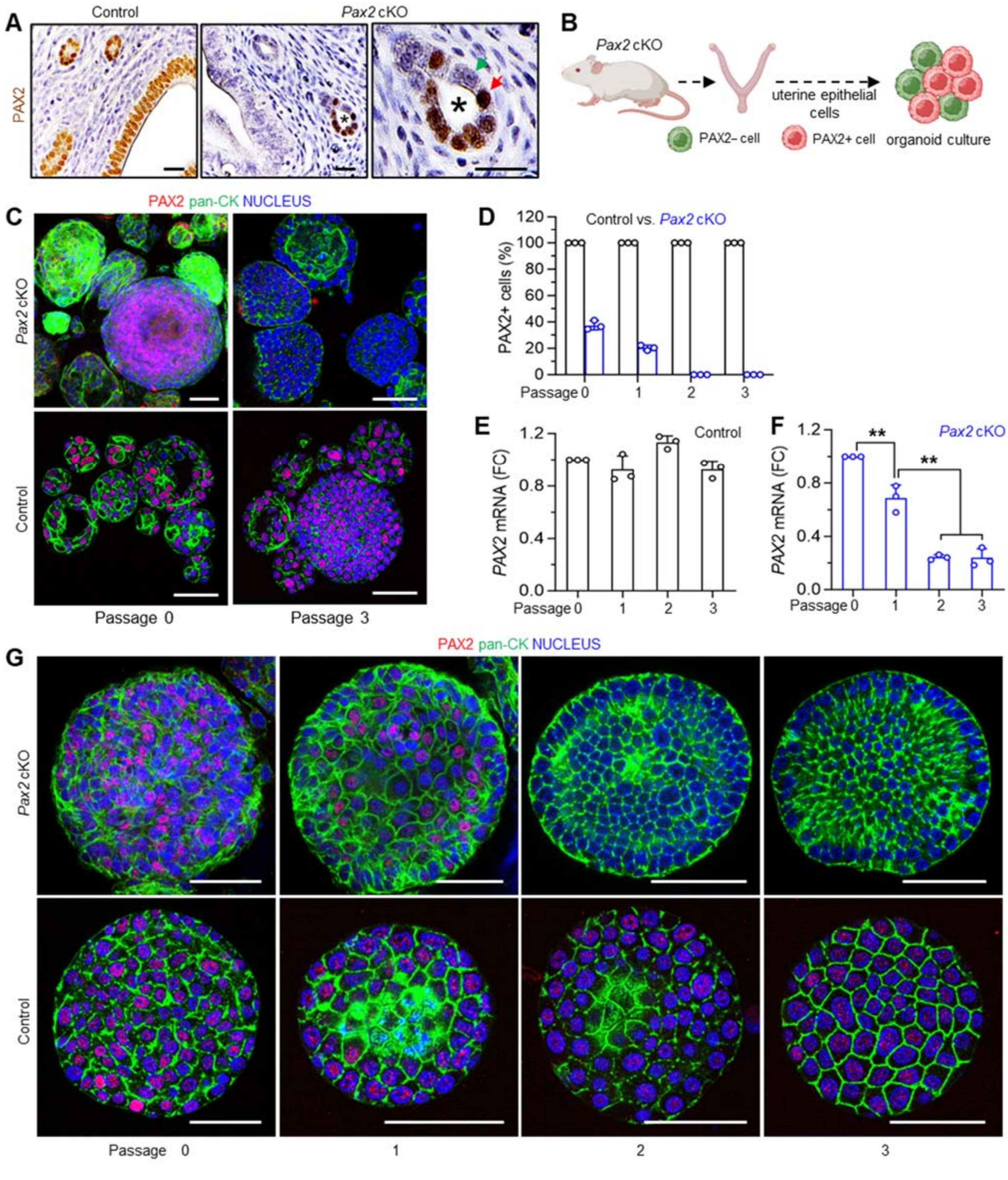

A
Control
Pax2 cKO
PAX2
B
Pax2 cKO
uterine epithelial cells
PAX2– cell
PAX2+ cell
organoid culture
C
PAX2 pan-CK NUCLEUS
Pax2 cKO
Control
Passage 0
Passage 3
D
Control vs. Pax2 cKO
PAX2+ cells (%)
Passage 0 1 2 3
E
Control
PAX2 mRNA (FC)
Passage 0 1 2 3
F
Pax2 cKO
PAX2 mRNA (FC)
**
**
Passage 0 1 2 3
G
PAX2 pan-CK NUCLEUS
Pax2 cKO
Control
Passage 0
1
2
3

**Supplementary Fig. 7. PAX2– cells outcompete PAX2+ cells in endometrial organoid culture.**

(**A**) PAX2 shows a mosaic pattern of PAX2 expression in *Pax2* mice at 8 weeks of age due to subtotal Cre-mediated recombination relative to control mice, in which PAX2 is expressed in all epithelial cells. An endometrial gland with mosaic PAX2 expression is indicated by an asterisk. The panel on the right shows this gland at higher magnification; the green arrow highlights epithelial cells lacking PAX2; red arrow highlights cells retaining PAX2. Bars=20 μm.

(**B**) Schematic of experimental approach of harvesting organoids from *Pax2*-mosaic mice (created with BioRender.com).

(**C**) Immunofluorescence staining for PAX2 and cytokeratin 8 (CK8) in control and *Pax2* organoids at passages 0 and 3 (n=3). Initially, PAX2+ cells were evident, which became depleted in *Pax2* organoid cultures after serial passaging. Bars=50 μm.

(**D**) Percentage of PAX2+ cells in control versus *Pax2* organoids across passages 0–3. Data represent mean±SEM (n=3).

(**E-F**) qRT-PCR of *Pax2* mRNA expression in control vs. *Pax2* organoids from passages 0 to 3. Data represent the mean±SEM (n=3); **P<0.01, unpaired 2-tailed t test.

**(G)** Confocal z-stack images of PAX2 and pan-cytokeratin (pan-CK) immunofluorescence in control and *Pax2* organoids from passages 0 to 3 (n=3). Bars=50 μm.
